# *Chamaeleo calyptratus* (veiled chameleon) chromosome-scale genome assembly and annotation provides insights into the evolution of reptiles and developmental mechanisms

**DOI:** 10.1101/2024.09.03.611012

**Authors:** Natalia A. Shylo, Andrew J Price, Sofia Robb, Richard Kupronis, Irán Andira Guzmán Méndez, Dustin DeGraffenreid, Tony Gamble, Paul A. Trainor

## Abstract

The family Chamaeleonidae comprises 228 species, boasting an extensive geographic spread and an array of evolutionary novelties and adaptations, but a paucity of genetic and molecular analyses. Veiled chameleon (*Chamaeleo calyptratus*) has emerged as a tractable research organism for the study of squamate early development and evolution. Here we report a chromosomal-level assembly and annotation of the veiled chameleon genome. We note a remarkable chromosomal conservation across squamates, but comparisons to more distant genomes reveal GC peaks correlating with ancestral chromosome fusion events. We subsequently identified the XX/XY region on chromosome 5, confirming environmental-independent sex determination in veiled chameleons. Furthermore, our analysis of the *Hox* gene family indicates that veiled chameleons possess the most complete set of 41 *Hox* genes, retained from an amniote ancestor. Lastly, the veiled chameleon genome has retained both ancestral paralogs of the *Nodal* gene, but is missing *Dand5* and several other genes, recently associated with the loss of motile cilia during the establishment of left-right patterning. Thus, a complete veiled chameleon genome provides opportunities for novel insights into the evolution of reptilian genomes and the molecular mechanisms driving phenotypic variation and ecological adaptation.

## INTRODUCTION

Squamates (lizards, snakes, and amphisbaenians), comprise over 11,600 species making it the largest order of reptiles^1^. Based on the fossil record, crown squamates originated about 200 million years ago^2^. The diversification of squamates is an extraordinary example of adaptive radiation, and whole-genome sequencing is instrumental in providing insights into their evolution, limited only by the small number of assembled squamate genomes^3,4^. Furthermore, studies of the early embryonic development of squamates have been hampered by the fact that, in the majority of egg-laying species, the embryos have completed key gastrulation and neurulation stages of development at the time of oviposition^5–7^. As a result, very little is known about early development events in reptiles such as the maintenance of pluripotency, gastrulation and left-right patterning, neurulation, neural crest specification and migration^5,8–12^. Hence there is a considerable need to sequence, assemble and annotate additional squamate genomes at chromosome scales to better understand the developmental mechanisms that drive phenotypic plasticity, diversity, and adaptation in concert with the evolution of complex organisms.

The family Chamaeleonidae comprises 14 genera and ∼228 species that exhibit a remarkable array of structural adaptations, including a projectile tongue, independently moveable turreted eyes, highly modified cranium, split hands and feet with differential syndactyly and zygodactyly, a prehensile tail, and rapid and complex color change^1,13^ Additionally, chameleon species exhibit an approximate 20-fold range in adult total length and a 2000-fold range in body mass.

Chameleons can be found distributed across Africa to the Middle East, in Madagascar, southern Europe, Asia and some islands in the Indian Ocean^14,15^ and they occupy an incredibly wide range of habitats including fynbos, forest, sandy desert, and grass^16–19^. Although most chameleons are arboreal, some species live predominantly on the ground^13^. Despite the wide geographical range and variety of ecosystems inhabited by chameleons, we currently have a poor understanding of the mechanisms underpinning their remarkable morphological adaptations, which have facilitated their radiation, survival, and reproduction in diverse environments. Collectively this makes chameleons an attractive model for studying the functional consequences of ecology, evolution, and development of phenotypic diversity.

In this study we have generated the assembly and annotation of the *Chamaeleo calyptratus* (veiled chameleon) genome at a chromosomal level. It is an emerging research organism for the study of early developmental processes and the diversification of evolutionary novelties^5^. In captivity veiled chameleons have well established husbandry and breed well, laying large clutches of eggs year-round at pre-gastrulation stages, in contrast to more established reptile research models^20–22^. Veiled chameleons use an XX/XY mode of sex determination, and here we identified the sex determining region on chromosome 5^23–25^. Our interest in morphogenesis lead us to examine *Hox* gene family, as well as the conservation of the left-right patterning Nodal cascade, revealing a unique pattern of gene conservation and loss, with a large potential to inform future evolutionary studies, given the position of reptiles in the phylogenetic tree.

The release of the veiled chameleon genome will facilitate functional genetic analyses through adoption of transgenesis and gene editing technologies developed in other reptiles^26,27^. Additionally, veiled chameleons are an invasive species in Florida and Hawaii in the USA, as well as in parts of Europe and Africa, and understanding their genetics and biology has the potential to aid management efforts^28,29^. Taken together with the recently published panther chameleon and dwarf chameleon genomes^30,31^, this also opens the door to a better understanding of genome evolution, evolutionary adaptation and early development in amniotes.

## RESULTS

### Genome sequencing and assembly

Prior to genome sequencing, we used cytometry-based methods to estimate the size of *Chamaeleo calyptratus* (veiled chameleon) genome to be 1.8Gb, which correlated well with the estimated average genome size of squamates^3,32^. We used male DNA for sequencing and genome assembly, resulting in a final total length of 1,803,547,962bp, which correlated with our cytological estimate^33^. Information about the *C. calyptratus* genome assembly is presented in Table 1.

**Table 1.**
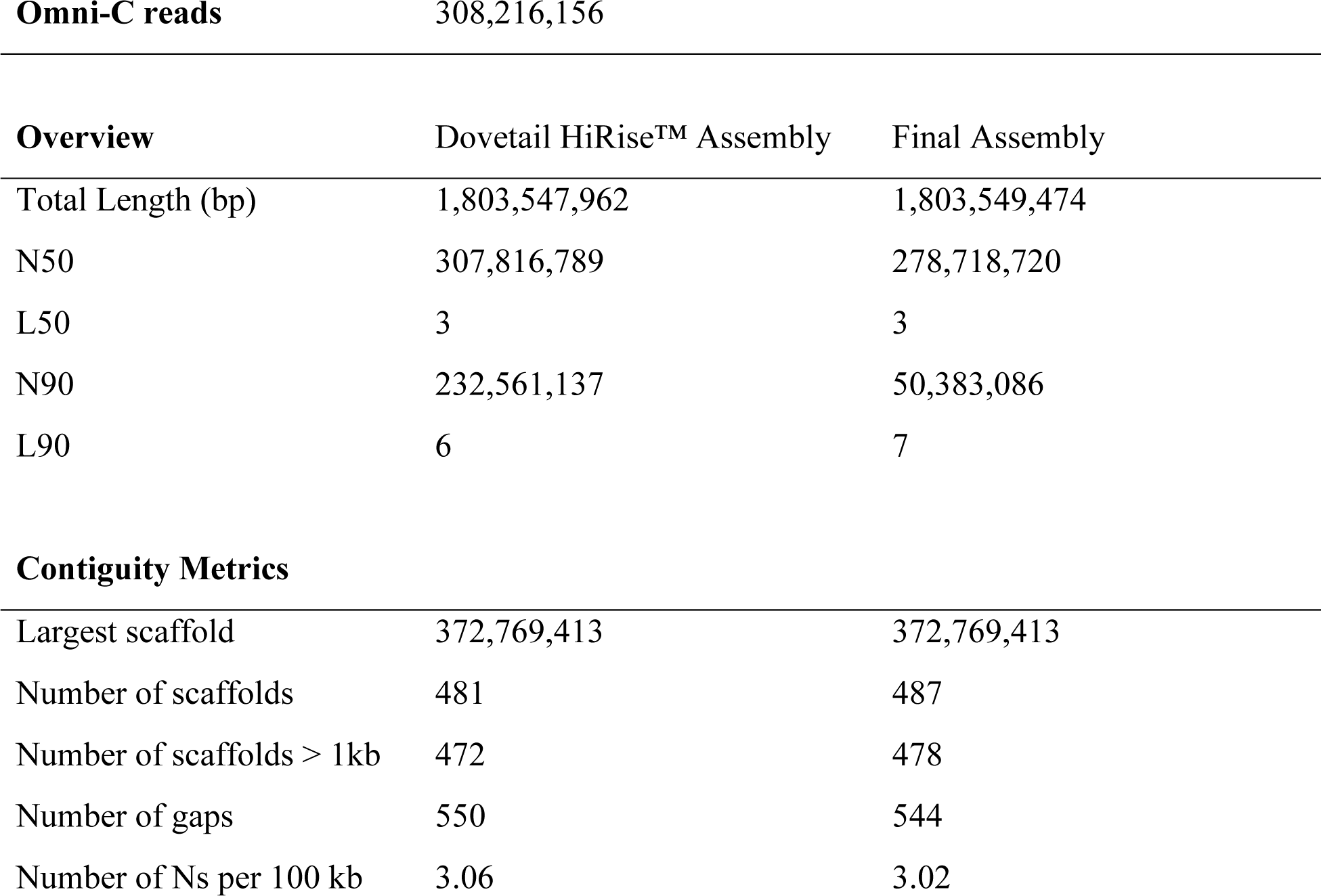
Assembly statistics for the *Chamaeleo calyptratus* genome.

### Chromosome anchoring

The veiled chameleon karyotype has been determined cytogenetically to be n=12 chromosomes, of which 6 are macrochromosomes, and 6 are microchromosomes^34,35^. Our initial genome assembly resulted in 481 scaffolds (Table 1), of which the six largest scaffolds contained 90% of the genetic information (L90) (Figure 1 A). To anchor assembled scaffolds to chromosomes, we used previously published sequencing results for individual flow sorted chromosomes^24^. Most of the reads, corresponding to individual macrochromosomes 1-5 mapped to individual scaffolds (Figure 1 B and Supplementary Figures S1-S6). However, a small percentage of reads from individual chromosomes mapped to different scaffolds, which likely represent repetitive elements (Figure 1 B and Supplementary Figures S1-S6). Surprisingly, reads from chromosomes 6-12 all mapped mostly to scaffold 3 (Figure 1 B and Supplementary Figure 3), and further investigation revealed clusters of reads, corresponding to individual chromosomes along scaffold 3 (Figure 1 C, Supplementary Figure S3).

**Figure 1.**
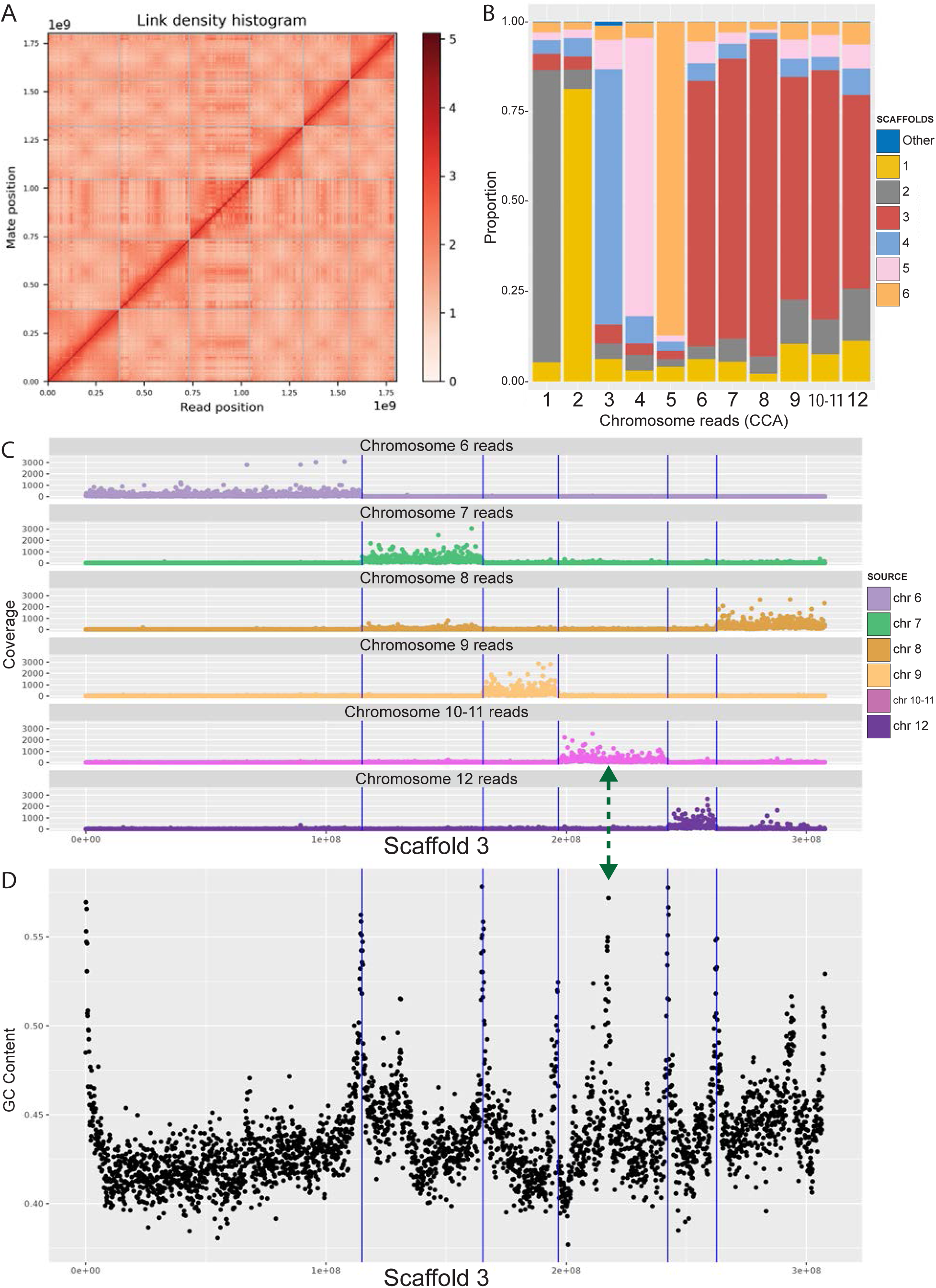
Anchoring scaffolds to karyotype. (A) Hi-C contact map. (B) Reads-scaffolds alignment. Reads from individual chromosomes (CCA 1-12) ^27^ were aligned against assembled scaffolds. The graph visually represents the proportion of reads from individual chromosomes, aligned to scaffolds. For each chromosome, the majority of the reads aligned to a unique scaffold. Most reads from chromosomes 6-12 are all aligned to scaffold 3. (C) Location of the chromosome-specific reads along scaffold 3. For visualization coverage was cut off at 3,000 reads. Full coverage maps are available in Supplementary Figure S3. (D) GC content along scaffold 3. High peaks of GC content correspond to chromosomal ends and align well with blocks of chromosome-specific reads in (C) (vertical lines). The green double-headed arrow indicates the junction point between chromosomes 10 and 11, which could not be separated for sequencing, and thus all reads binned together in (C).

To manually break up scaffold 3 into individual microchromosomes, we relied on the location of the reads from individual flow-sorted chromosomes^24^, synteny analysis across other species, as well as GC content along the scaffold. Chromosome edges had higher GC content, highlighting the presumed ends of chromosomes (Figure 1 D, Figure 2, Supplementary Figure S8). We further identified gaps, containing 100 Ns, marking the locations of computational scaffold fusions. Although chromosomes 10 and 11 were not resolved through flow sorting^24^, we identified a high GC content peak that bisected the 10-11 sequence region, coupled with a gap in the assembly (Figure 1 C, D green arrow). We labeled the two resulting scaffolds chromosomes 10 and 11 in descending scaffold size order.

**Figure 2.**
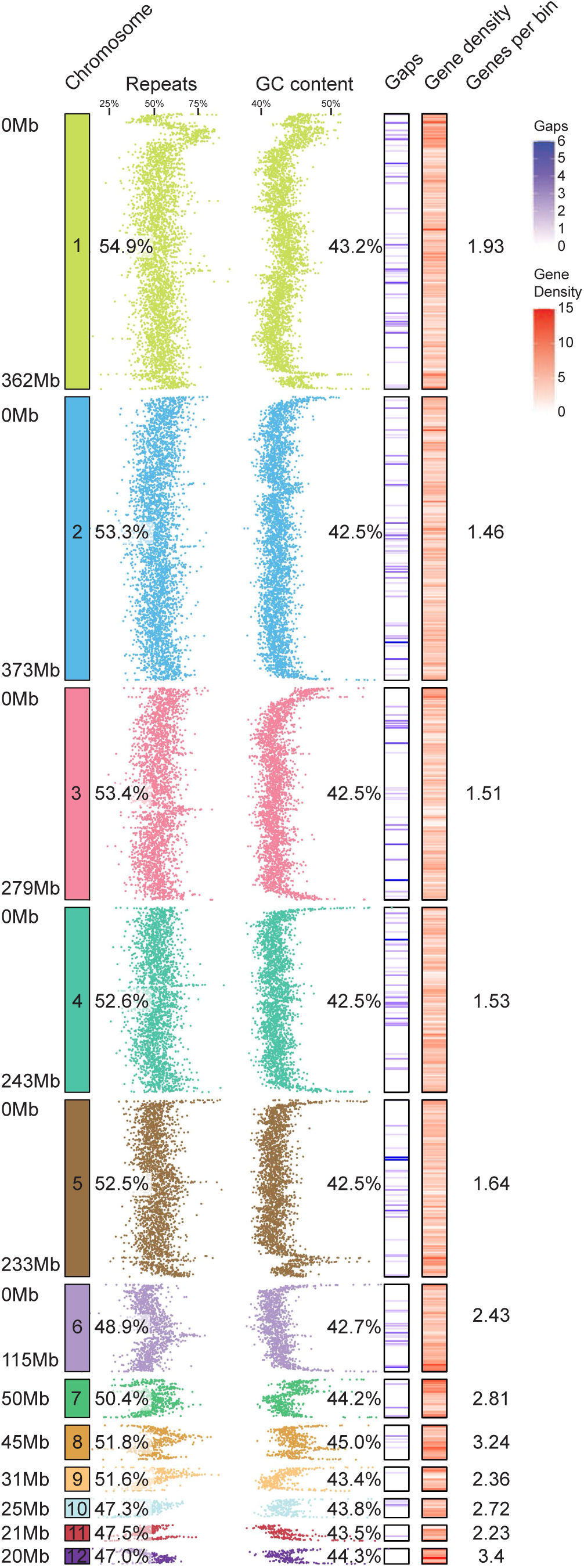
Structure and content of the veiled chameleon genome. 12 to-scale chromosomes, with individual scaffold sizes indicated on the left, rounded to the nearest Mb. The schematics reveal regional variation in genomic repeat content, GC content, gaps in the assembly (541 gaps across 12 largest scaffolds), and gene density, as determined for individual 100kb bins. The numerical values represent % repeat content, % GC content and the average number of genes in every 100kb bin. The microchromosomes (7-12) have significantly lower amount of repetitive DNA (p≤0.0219), significantly higher GC content (p≤0.0004) and significantly higher gene density per 100kb bin (p≤0.0016), as determined by unpaired two-sided t-test. Mb-megabases.

Another contradiction between our original assembly and individual chromosome sequencing results involved scaffolds 1 and 2 (arranged by length) and reads for flow-sorted chromosomes 1 and 2 (CCA1 and CCA2)^24^. Reads from CCA1 mapped to scaffold 2 and reads from CCA2 mapped to scaffold 1 (Figure 1 B, Supplementary Figures S1, S2). Previous reports have identified a 45S rDNA fluorescent *in situ* hybridization signal on chromosome pair 1, and interstitial telomeric repeats in the pericentromeric region of the largest metacentric chromosome pair^34,36^. These data were consistent with the sequencing results of CCA1^24^. In our assembled genome 45S rDNA as well as the highest enrichment of interstitial telomeric repeats, map to scaffold 2 (Supplementary Figure S9, Supplementary Table 1). rDNA is highly repetitive and poses challenges for accurate genome assembly, frequently collapsing the region^37^. Thus, we conclude that scaffold 2 corresponds to chromosome 1, but appears shorter due to insufficient resolution of the rDNA region. The final scaffold-chromosome nomenclature between different studies is outlined in Supplementary Table 1, and the relationship between originally assembled scaffolds and finalized 12 chromosomes is outlined in Supplementary Figure S7.

### Genome characteristics and annotation

The final assembly of the veiled chameleon genome consists of 487 scaffolds (Table 1), but for the remainder of the analysis we will focus on the 12 largest scaffolds, representing the 12 chromosomes in veiled chameleon. The 12 assembled scaffolds range in size from ∼373Mb to ∼20Mb, and currently still have 544 gaps in the assembly (Table 1, Figure 2).

The density of repetitive elements is highly variable across the chromosomes, with some chromosomes having higher repeat density near the termini, while others have lower density. Overall, microchromosomes 7-12 have significantly lower mean repeat density than macrochromosomes 1-6 (Figure 2). This observation is consistent with previous reports of lower average repeat content on microchromosomes in rattlesnake and chicken^38^. GC content is also variable across chromosomes, with high levels on chromosome ends, and multiple intrachromosomal peaks (Figure 2, Supplementary Figure S8). Overall, mean GC content is higher on microchromosomes than macrochromosomes.

We hypothesized that since the chromosomal ends have high GC content, the interstitial peaks of GC content may be the result of ancestral chromosomal fusions. Thus, we interrogated the distribution of interstitial telomeric repeats (TTAGGG)n across the genome (Supplementary Figure S9). Our results revealed a high repeat density in centromeric region of chromosome 1 (Supplementary Figure S9), consistent with a prior publication^36^. Nevertheless, this single enrichment correlated poorly with numerous interstitial peaks of GC content across the genome (Supplementary Figure S8). Ultimately, we scanned the genome for 6-mers which had positive correlation with GC content peaks and the top 10 sequences with positive (Supplementary Figure S10, Panel 1), negative (Supplementary Figure S10, Panel 2) or no correlation (Supplementary Figure S10, Panel 3) are presented in Supplementary Table 2.

We used Helixer to predict gene models using only the genome assembly, and further supplemented these with previously published mRNA IsoSeq^12,39,40^. The resulting structural annotations produced higher BUSCO scores^41^ and took less time to generate (8 hrs) than other commonly used tools like MAKER^42^. The final BUSCO scores of genome completeness, using Metazoa, vertebrate, Tetrapoda and Sauropsida datasets for genome and protein predictions are available in Supplementary Table 3. The gene density analysis revealed higher average gene density on microchromosomes, compared to macrochromosomes (Figure 2).

### Synteny analysis

As more reptilian genomes are assembled, and scaffolds are anchored to chromosomes, the evaluation of chromosomal reshuffling between species and studies of genomic evolution become increasingly possible^3^. Thus, we carried out pairwise synteny analysis between veiled chameleon, dwarf chameleons *(Bradypodion ventrale* and *Bradypodion pumilum*)^31^, panther chameleon^30^, brown anole^43^ and chicken genomes.

Comparative syntenic maps between veiled chameleon and other reptiles revealed remarkable chromosomal conservation among squamates examined. Macrochromosomes 3-6 in veiled chameleon correspond 1:1 with chromosomes 3-6 in the brown anole (Figure 3, Supplementary Figure S11 Panel 4). Furthermore, chromosomes 7, 9, 10 and 11 also have counterparts in brown anole genome, and chromosomal reshufflings resulted in fusions of large syntenic blocks (Figure 3, Supplementary Figure S11 Panel 4). Likewise, dwarf chameleon genomes exhibit similar architecture to both brown anole and veiled chameleon genomes (Figure 3, Supplementary Figure S11 Panels 2, 3). Remarkably, we found high degree of chromosomal reshuffling in the panther chameleon genome, which was particularly notable for panther chameleon chromosome 1, which is syntenic to veiled chameleon chromosomes 2, 3 and 8 (Figure 3, Supplementary Figure S11 Panel 1).

**Figure 3.**
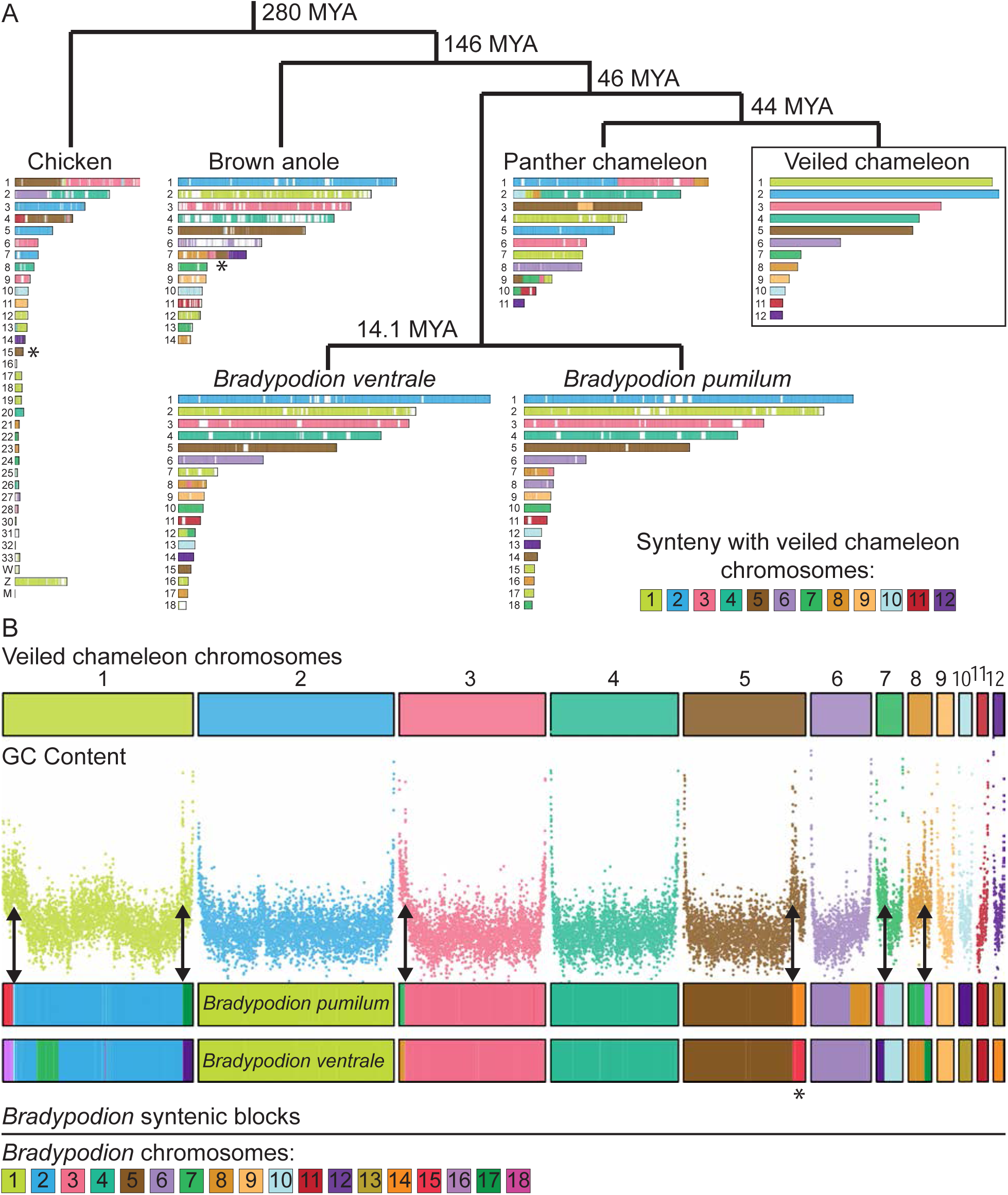
Synteny analysis. (A) Synteny between veiled chameleon, panther chameleon, dwarf chameleons *Bradypodion ventrale* and *Bradypodion pumilum*, brown anole, and chicken genomes. Phylogenetic distance determined using TimeTree database^100^. Chromosomes are painted according to synteny with veiled chameleon chromosomes. Reciprocal comparisons between veiled chameleon and individual species are available in Supplementary Figure S11. (B) Synteny analysis between veiled chameleon and dwarf chameleons (*Bradypodion*), with comparison to regional GC content. Chromosomes are painted according to synteny with dwarf chameleon chromosomes. Double-headed arrows indicate regions where fusions of dwarf chameleon syntenic blocks correlate with regions of high GC content in veiled chameleon genome. Asterisks denote the location of a genomic block, syntenic with chromosome 15 in chicken, previously identified as a conserved X-linked sex chromosome element in all pleurodont iguanas (except basilisks and their relatives; chromosome 5 in chameleon, chromosome 7 in brown anole)^126^.

A comparison of syntenic blocks between dwarf chameleons (*Bradypodion*) and the veiled chameleon genome, alongside GC content across the veiled chameleon genome revealed that GC-content peaks match up with the borders of syntenic blocks (Figure 3 B arrows). Given that chromosome ends in chameleons contain high levels of GC content, these intrachromosomal GC peaks likely represent the locations of ancestral chromosome fusion events, supporting our prior hypothesis. Furthermore, exactly 6 regions show highest GC peaks and synteny to dwarf chameleon microchromosomes. Dwarf chameleons are thought to have the ancestral chameleon karyotype (n=18)^44^, whereas veiled chameleon karyotype has been reduced to n=12 chromosomes. Therefore, we think the 6 identified syntenic regions represent the additional ancestral microchromosomes, thus allowing us to identify all 6 ancestral microchromosomes in the veiled chameleon genome.

Additional comparison to a more distantly related chicken genome revealed other regions where syntenic blocks align with peaks in GC content both on macrochromosomes and microchromosomes (Supplementary Figure S12 arrows). We hypothesize that higher peaks representing more recent fusion events, and lower peaks more ancient events.

### Genetics of sex determination in veiled chameleons

We have previously shown that veiled chameleons utilize XX/XY genetic sex determination^23,24,45^. However, the chromosomal sex determination region has remained elusive^24,34,35^. More recently, sex-specific genetic markers identified chromosome 5 as the sex chromosome in veiled chameleons^23,24^. For genome assembly and annotation, we sequenced male gDNA to ensure coverage of both the X and Y chromosomes, expecting only half the coverage of autosomes for the sex chromosomes. Contrary to our prediction, all scaffolds exhibit uniform sequence coverage, including chromosome 5. This is consistent with sex chromosomes that are newly evolved and poorly differentiated from each other^46^.

Recently evolved, homomorphic sex chromosomes can be detected through the identification of sex-specific genetic markers from restriction-site associated DNA sequencing (RAD-seq) data^47^. The male/female F_ST_ value then further defines the non-recombining region of the sex chromosomes^47,48^. Our analysis of male/female F_ST_ across veiled chameleon chromosomes revealed a defined peak at the terminal part of chromosome 5 (Figure 4 A arrow, B). Notably, all 13 male-specific markers mapped within the most terminal 11.2Mb of chromosome 5, with 11 of them tightly mapped to the most terminal 0.85Mb, overlapping the F_ST_ peak (Figure 4 A, B, Supplementary Table 4). Only a single gap in the assembled genome maps to this region, located at Chr5:872,188bp (Figure 4 C).

**Figure 4.**
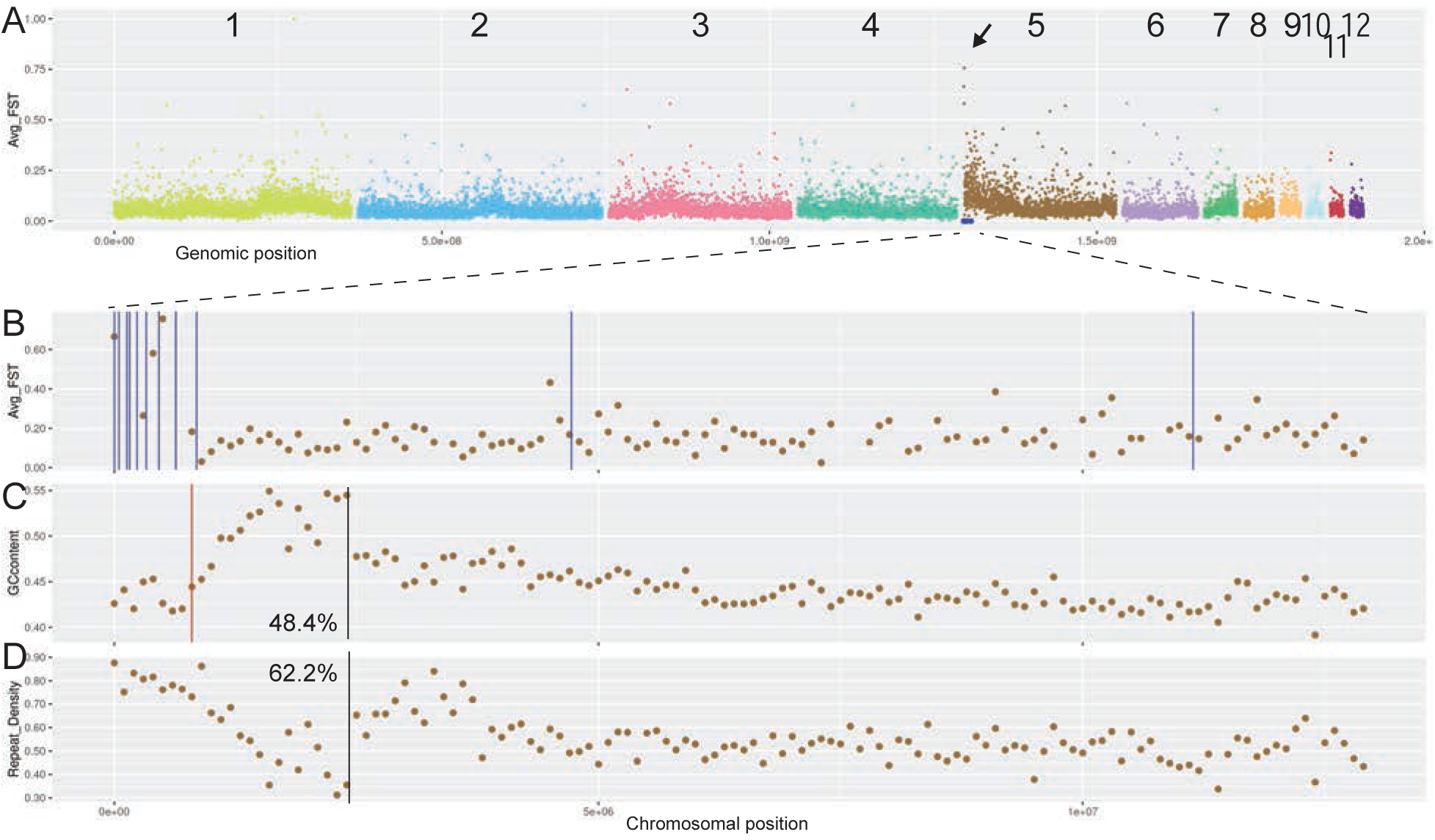
Sex determination in veiled chameleons. (A) RAD-seq M/F F_ST_ scan in 100kb bins. Arrow points to a peak in M/F F_ST._ Blue dots denote the location of 13 unique male-specific markers, as previously identified ^33^. (B) RAD-seq M/F F_ST_ scan in 100kb bins, zoomed in on the first 13 Mb of chromosome 5. Vertical blue lines denote the location of unique male-specific markers ^33^. Exact genomic coordinates of marker locations available in Supplementary Table 4 (C) GC-content in the zoomed-in region. The red vertical line denotes the single gap in the genome assembly in this region. Black vertical line denotes the approximate inflection point of GC content. 48.4% - GC content in the region leading up to the black line. (D) Repeat content in the zoomed-in region. Black vertical line denotes the approximate inflection point of content levels. 62.2% - portion of repetitive elements up to the black line.

The GC content in the terminal region of chromosome 5 peaks around 2Mb (Figure 4 C, D, black vertical lines), which sets it apart from other chromosomes with elevated GC content at the chromosomal edges (Supplementary Figure S8). We hypothesize that the sex determination region in veiled chameleons arose as a result of an inversion of the most terminal 2Mb, resulting in the GC peak in the location of re-fusion. The current assembly does not resolve the differences between the X and Y chromosomes in this region, thus it is unknown which of the sexes may harbor this inversion. Recombination is often prevented on sex chromosomes as a result of an inversion, and this results in the accumulation of transposable elements. Consistent with that scenario, the repeat density is particularly high in this region of chromosome 5, at 62.2%, compared to 52.4% for the rest of chromosome 5 (Figure 4 D).

Next, we identified 188 genes, spanning the 13 male-specific markers (Chr5: 1-11,324,268) (Supplementary Table 5). Unfortunately, none of these are known sex determination genes. Our annotation predicted only 3 genes (*Lrwd1*, *CCA1g016450000.1* and *CCA1g016449000.1*) in the area that overlaps with the first 11 markers (Chr5:1-825,560), and of those, *Lrwd1* (Permanent gene ID available in Supplementary Tables 5 and 6) is involved in human spermatogenesis^49^ and Xi silencing^50^. Improved mRNA-seq-based annotation will likely reveal additional genes in this region. Overall, we identified 28/188 genes with known links to sex determination and reproduction (Supplementary Table 5). Among them, *TNPO3* and *UBE2H* stand out as Y-linked genes in skinks^51,52^. We then mapped the skink Y-linked transcripts onto the veiled chameleon genome and identified 13 markers that overlap the ∼11.2Mb sex determination region (non-identical to chameleon Y-specific markers)^23,51^. Interestingly, the opposite end of chromosome 5 is syntenic with chicken chromosome 15, a conserved X-linked sex determination region across pleurodont iguanas (Figure 3, Supplementary Figure S12 A, asterisks)^53–61^.

### Environmental impact on sex determination in veiled chameleons

Although we have identified sex chromosomes in the veiled chameleon, environmental factors may also play a role in veiled chameleon sex determination^45,62,63^. Therefore, we incubated eggs at 24, 26, 28 and 30⁰C to detect any changes in sex ratios^45^. After harvesting embryos we used morphological characteristics to determine their sex phenotypically (Supplementary Figure S13 A-D), and a sample of DNA was then used to determine their genotypic sex^23^.

We observed no significant difference in sex ratio distribution across all temperatures (Supplementary Figure S13 E). At 28⁰C and 30⁰C five individuals were phenotypically identified as males but were genetically females. These were most likely mis-identified as phenotypic males, since the skin fold on female hindlimbs can occasionally be large enough to be mistaken for a bony heel spur (Supplementary Figure S13 B, D).

We also tested whether there was any correlation between the sex and the weight of an egg. Egg size and weight varied from clutch to clutch, with some variation within clutches. Mean egg weight however was similar between the sexes at 24, 26, and 30⁰C, and standard deviation values were high for all temperatures (Supplementary Figure S13 F). We only found a significant difference in weight between male and female eggs at 28⁰C, with male eggs being heavier.

Although this observation at 28⁰C is similar to another report^63^, it is contrary to observations in other studies^45,63^. Therefore, we believe it may be a chance event, as the overwhelming volume of data across all temperatures found no link between egg size or weight and sex. We also noted fluctuations in weight, based on temperature, which are likely due to environmental factors (Supplementary Figure S13 F). Over the course of development, egg weight typically increases, and this was reflected in our data, with a mean increase of 12.08% for female eggs, and 13.54% for male. However, the difference between the two sexes was not statistically significant (Supplementary Figure S13 G). Therefore, we conclude that veiled chameleon sex is genetically determined with temperature and egg weight changes having no effect on or correlation with sex determination.

### Nodal left-right patterning cascade

Left-right patterning in reptiles is motile-cilia independent and has been receiving renewed attention in the field, albeit with limited analysis of gene conservation^10,12,64,65^. The analysis of left-right patterning in reptiles provides a key evolutionary link to understand the nuances of the process in amniotes, including humans. Recently, we identified two *Nodal* transcripts (Supplementary Table 6) in veiled chameleons with unique patterns of expression during gastrulation and left-right patterning^12^. Sequence analysis suggested that these two transcripts are orthologs of *Nodal1* and *Nodal2*, which were duplicated in the gnathostome lineage^10,12,66^.

Analysis of the annotated veiled chameleon genome revealed that the *Nodal* transcripts indeed derived from two orthologous genes – *Nodal1* and *Nodal2* – and both genomic regions revealed considerable conservation of syntenic groups (Figure 5 A, B).

**Figure 5.**
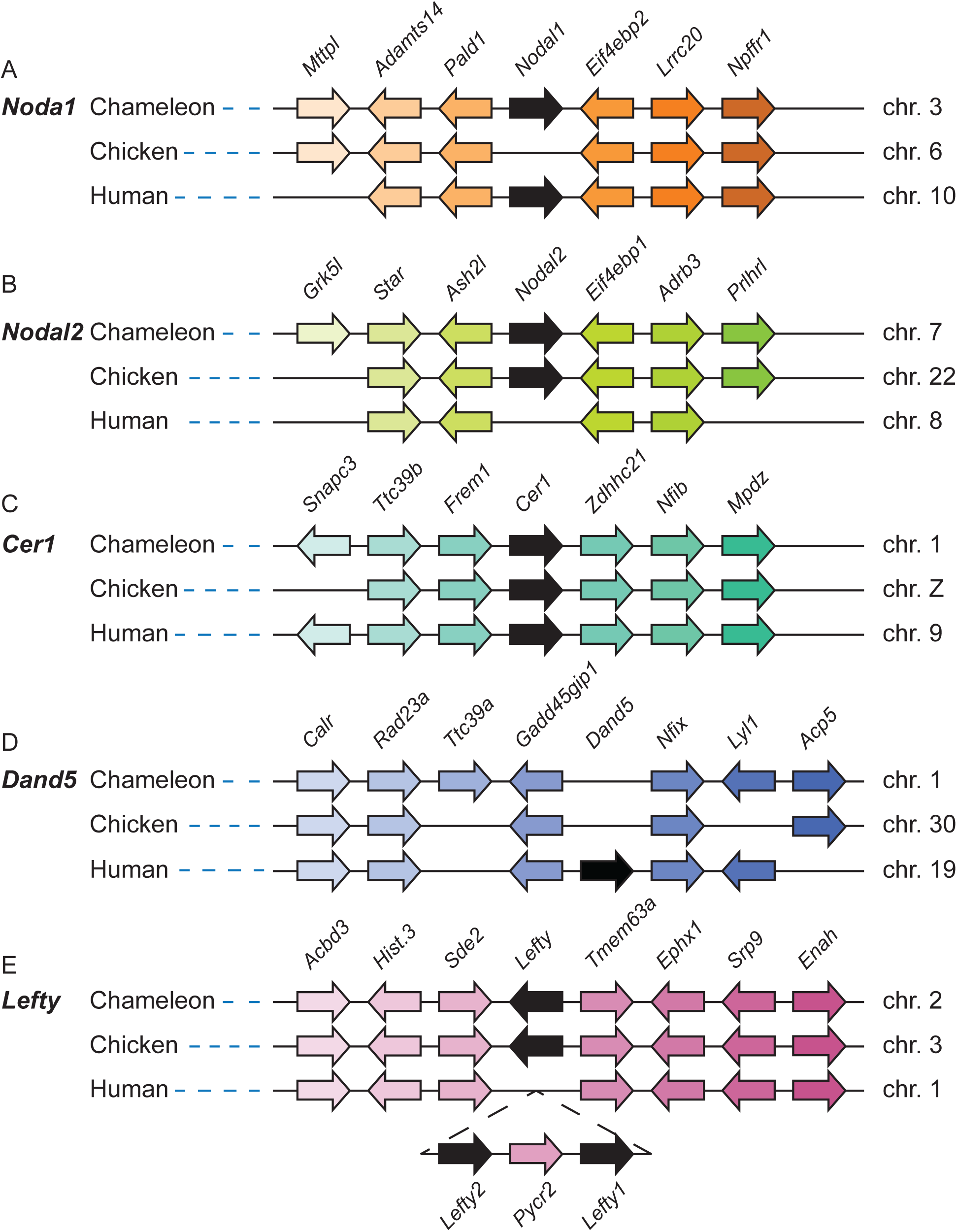
Nodal left-right patterning cascade gene linkage analysis. (A) Linkage analysis of the *Nodal1* gene region in veiled chameleon, chicken and human genomes shows absence of *Nodal1* in the chicken genome. (B) Linkage analysis of the *Nodal2* gene region between veiled chameleon, chicken and human genomes shows absence of *Nodal2* in the human genome. (C) Linkage analysis of the *Cer1* gene region between veiled chameleon, chicken, and human genomes. (D) Linkage analysis of the *Dand5* gene region between veiled chameleon, chicken and human genomes shows absence of *Dand5* in veiled chameleon and chicken genomes. (E) Linkage analysis of the *Lefty* gene region between veiled chameleon, chicken and human genomes shows presence of only one *Lefty* gene in chameleon and chicken genomes, and gene duplication in the human genome. Black arrows denote the genes of interest. Gene orientation is indicated by the direction of the arrows. Orthologous genes are colored with the same color.

We then evaluated genomic conservation of additional members of the Nodal left-right signaling cascade. We had previously identified *Cer1*, a Dan family member, as the repressor of Nodal in the lateral plate mesoderm of neurulation stage embryos^12^. Our analysis confirmed the presence of a single *Cer1* gene in the veiled chameleon genome (Figure 5 C). In contrast, another Dan family member, *Dand5*, was lost from the chameleon genome (Figure 5 D), as has been reported in chickens, geckos, and turtles, thus providing further support regarding its absence in all reptilian genomes^10,12,64^. The syntenic region including surrounding genes is, however, well conserved (Figure 5 D).

Lefty is another repressor of Nodal signaling that acts to restrict *Nodal* expression to the left side of the embryo. Similar to chickens, veiled chameleons have a single *Lefty* gene, in contrast to mammals and fish with duplicated *Lefty* genes (Figure 5 E)^65^. Pitx2 is a transcription factor downstream in the Nodal cascade, and we have confirmed the conservation of a single *Pitx2* gene in the veiled chameleon genome.

Recently a group of genes was identified in the context of left-right patterning that have been lost from the published reptilian genomes^64^. The five genes include *Dand5*, *Pkd1l1*, *Mmp21*, *Cirop* and *C1orf127*. It is currently unknown what functional relationship these genes may have in the L-R patterning cascade. We analyzed the genomic regions containing these genes and confirmed their absence from the veiled chameleon genome.

### *Hox* genes in veiled chameleons

*Hox* genes comprise a large family homeobox transcription factors, that specify the patterning and development of the body plan of an embryo, and reptiles are well known for their extreme skeletal modifications, like the carapace in turtles and the absence of limbs in snakes^67^. Modifications of *Hox* gene expression are therefore central to body plan diversification. The ancestral tetrapod is believed to have had 41 *Hox* genes, arranged in four clusters in the genome, after two rounds of ancestral duplication events^68^. However, turtles, crocodiles, birds and placental mammals have 39 *Hox* genes, having lost *HoxC1* and *HoxC3*.

Our analysis of *Hox* gene clusters in the veiled chameleon genome revealed four clusters of genes. *HoxA* and *HoxB* clusters are both located on chromosome 6, the *HoxC* cluster is on chromosome 1, and the *HoxD* cluster is on chromosome 2 (Figure 6). Due to the similarity of *Hox* genes, we manually curated many gene annotations in the four clusters. Although *HoxC1* has been repeatedly lost in amniotes and is only found in some lizards^69^, we identified *HoxC1* in veiled chameleon and have detected evidence of its expression in our mRNA sequencing samples. Likewise, we were able to identify *HoxC3* and evidence of its expression in veiled chameleons. Thus, veiled chameleon possesses the full tetrapod set of 41 *Hox* genes.

**Figure 6.**
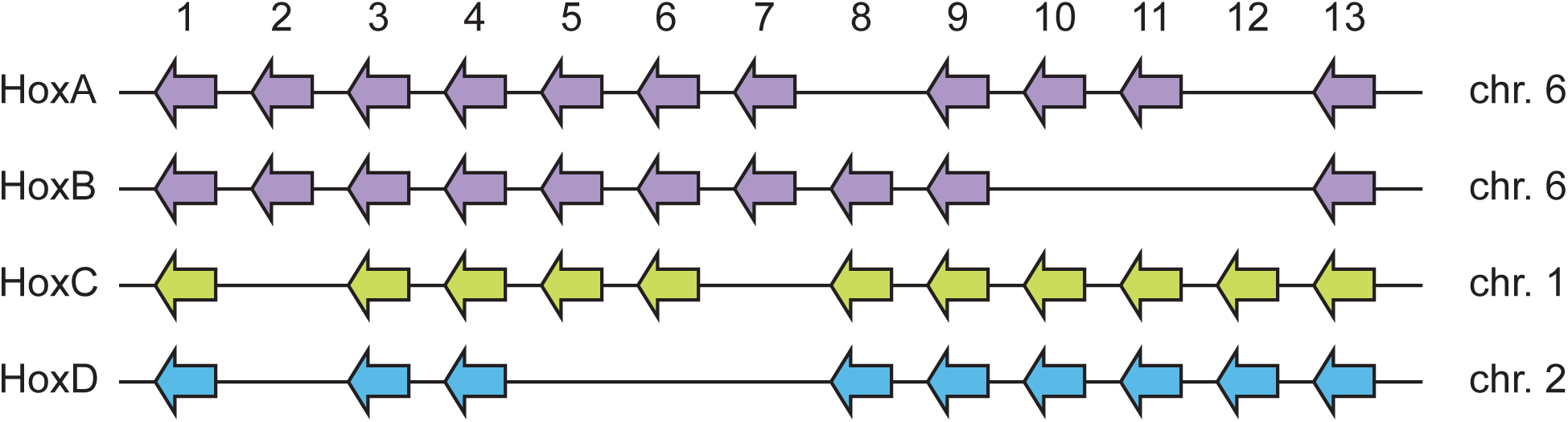
Hox gene clusters in veiled chameleon genome. Veiled chameleon genome contains four clusters of Hox genes, located on chromosomes 6 (two clusters, purple), 1 (green) and 2 (blue), for a total of 41 genes. Genes are colored to match the chromosome identity.

## DISCUSSION

Chamaeleonidae comprises an estimated 228 species, which are characterized by evolutionary morphological and behavioral adaptations, which include a cranial casque, forelimb and hindlimb syndactyly and zygodactyly, a projectile tongue, prehensile tail and the ability to color change ^1,5,70–74^. Despite the wide geographical range and variety of ecosystems inhabited by chameleons, we currently have a poor understanding of the mechanisms underpinning their remarkable morphological adaptations, which have facilitated their radiation, survival, and reproduction in diverse environments. Chameleons are therefore an ideal model for studying the genetic events that regulate the evolution of morphological adaptation.

In this study, we report the chromosomal-level assembly and annotation of the *Chamaeleo calyptratus* (veiled chameleon) genome^3,30^. This new reference genome provides a foundation for molecular, genetic, quantitative, and population genomic studies, and a resource for accelerating research into the evolution of adaptive traits in squamates to complement the advances brought about by sequencing of other reptiles.

The recent explosion of newly sequenced high-quality reptile genomes has ushered in a new era of comparative genomics and two of the most active areas of genomic research in reptiles, and squamates in particular, involve microchromosome evolution and the evolution of sex determination^75,76^. Microchromosomes are lost in mammals, but are otherwise common across the animal kingdom, with a huge variation in numbers in some reptiles, with multiple fusions and reshufflings between species^77^. Although small, they have high gene densities, higher GC content than macrochromosomes, low numbers of transposable and repetitive elements, high rates of recombination, and house many rapidly evolving genes, like the genes for venom production in snakes ^38,77,78^. In the nucleus, microchromosomes congregate in the central nuclear territory with extensive inter-chromosomal interactions^78,79^. This biological behavior of microchromosomes, as shown here, has made them challenging to resolve by Hi-C-based methods of genome assembly^38,43,80^.

Veiled chameleon chromosomes 7-12 have traditionally been classified as microchromosomes, based on their significant difference in size, compared to macrochromosomes 1-6^34,35^. In our assembly chromosomes 6-12 were computationally fused into scaffold 3, hinting at tight inter-chromosomal interactions between them. Despite being ∼115Mb in size, chromosome 6 has many features of microchromosomes. The average GC content, repeat and gene density on chromosome 6 are more similar to microchromosomes than macrochromosomes 1-5 (Figure 2). These observations suggest that microchromosomes deserve a functional definition, and more targeted research to understand what drives their nuclear territory localization, genetic composition, frequent fusions, and rearrangements.

With respect to the evolution of sex determination in reptiles, squamates in particular exhibit numerous transitions involving XX/XY, ZZ/ZW and environmentally dependent sex determining systems, with some transitions occurring among closely related species^47,75,81,82^. Analysis of diverse sex determination regions has revealed similar chromosomal regions being used across clades, hinting at the potential existence of an ancestral sex chromosome, or an equally likely scenario that some genes and genetic regions are more prone to take on the roles of master regulators of sex determination, evolving into their capacity through gene duplication and mutagenesis^83–86^.

We identified the terminal region of chromosome 5 as the sex determination region in veiled chameleon. It is a remarkably small region, with most male-specific markers mapping to the first ∼0.85Mb of the scaffold, and all 13 markers mapping within the first ∼11.2Mb^23^. The 11.2Mb region contains 188 genes, including transcription factors and many genes of unknown function. A prior genetic analysis of the male-specific markers confirmed their specificity in *Chamaeleo chamaeleon*^23,81^, indicating that this sex determination region is conserved in the genus *Chamaeleo*, placing its origin at ∼11.7 million years ago^87^. In contrast, the Malagasy giant chameleon (*Furcifer oustaleti*) utilizes ZZ/ZW mode of sex determination, and the panther chameleon (*Furcifer pardalis*) exhibits a Z_1_Z_1_Z_2_Z_2_/Z_1_Z_2_W system of multiple sex chromosomes^82^. The mode of sex determination is currently unknown for dwarf chameleons, despite the availability of two newly sequenced genomes^31^.

We noted that part of the sex determination region in veiled chameleon also overlaps with the Y-linked region in skinks^51,52^, supporting the observation that many sex determination regions across reptiles have the same ancestral genomic origin^84,86^. More work therefore is necessary to further delineate the sex determination region in veiled chameleon. A male chameleon, the heterogametic sex, was used for genome assembly in this study, and at current resolution we were not able to clearly differentiate the X and Y sequences. It is possible that due to their similarity, the X and Y chromosomal regions may have been computationally intercalated in our assembly. However, it is unclear if X and Y chromosomes are of equal length and have a highly divergent sex determination region, or whether the X chromosome is missing the Y-linked region. We speculate that the sex determination region may have arisen as a result of an inversion at the beginning of chromosome 5, based on the presence of a GC peak around Chr5:2Mb .

Curious about the evolutionary origin of this region of high GC content, we mapped the GC content across all veiled chameleon chromosomes and noticed a pattern of multiple scattered peaks (Figure 2, 3, Supplementary Figure S8). GC isochores are well known in genomes, although their evolutionary origin has remained a mystery. When we coupled that analysis with synteny analysis of several genomes, it revealed a remarkable pattern, strongly suggesting that the GC peaks across chromosomes are likely remnants of ancestral chromosome fusions (Figure 3, Supplementary Figure S12). A comparison of GC content and syntenic blocks between the veiled chameleon and dwarf chameleon genomes makes it possible to identify the locations of ancestral fusions of 6 microchromosomes. Dwarf chameleons are thought to have the ancestral chameleon karyotype (n=18)^44^. Therefore, we can identify all 12 present and ancestral microchromosomes in the veiled chameleon genome.

We are not aware of other reports that use GC content and synteny analysis to understand the process of chromosome evolution. We hypothesize that higher GC peaks represent more recent fusion events, and lower peaks represent evolutionarily older events. It is conceivable that as more chromosome-level assemblies of reptile genomes emerge, the patterns of GC content and microchromosome rearrangements may be used to trace the evolution of chromosomes, supplementing current cytological and synteny analyses to ultimately determine the karyotypes of the ancestral genomes^44^. It should also be noted that regions of elevated GC content are often recombination hotspots, and this property of the chameleon genome will need to be investigated in the future.

In summary veiled chameleons are an ideal research organism for the study of evolutionary and developmental processes in non-avian reptiles due to their ease of husbandry, and pre-gastrulation stage of development at the time of oviposition^5,12,20,21,71,72^. Aided by our newly sequenced and annotated veiled chameleon genome, preliminary analyses of gene conservation have already revealed unique aspects of veiled chameleon development. Veiled chameleons possess 41 *Hox* genes, the most extensive set in amniotes, an evolutionary feature that likely contributes to their unique body plan^69^. The analysis of the genes in the left-right patterning cascade revealed two *Nodal* genes, and loss of the Nodal inhibitor *Dand5*, as well as *Pkd1l1*, *Cirop, Mmp21* and *C1orf127*^64^. Loss of the same genes from reptilian genomes and the genomes of even-toed ungulates, none of which have left-right organizer possessing motile cilia, suggest the involvement of these genes in left-right patterning, with likely functions in signal titration and relay.

Rapid advances in genomic technologies and approaches will continue to further our understanding of the molecular drivers of phenotypic plasticity and well-annotated genomes will facilitate the application of gene editing to study the contributions of individual genes to the evolution of early developmental processes, like gastrulation, left-right patterning, axial body patterning, neurulation and organogenesis. Our high quality chromosome-level veiled chameleon genome will serve as a reference for future research on chameleon, reptile, and amniote genomic and morphological evolution. Chameleons exhibit many unique characteristics providing biologists with numerous opportunities to study novel evolutionary innovations and processes.

Given the incredible diversity within chameleons and their specialized behavior, they are an attractive group for studying the functional consequences of phenotypic diversity.

## METHODS

### Animal husbandry

All animal experiments were conducted in accordance with the Stowers Institute for Medical Research Institutional Animal Care and Use Committee approved protocol 2020-115. Veiled chameleon husbandry was performed in our Reptiles and Aquatics Facility as described previously ^20,21,71^ following the protocols which are publicly available here: dx.doi.org/10.17504/protocols.io.bzhsp36e.

### Data availability

All raw data and assembled genomes are available via the National Center for Biotechnology Information under the accession PRJNA1106902. Veiled chameleon genome, as well as all previously published RNA datasets can be downloaded, browsed and searched on publicly available browser at simrbase.stowers.org.

All original data underlying this manuscript is available and can be accessed from the Stowers Original Data Repository at http://www.stowers.org/research/publications/LIBPB-XXXX.

### Tissue collection

For DNA collection, two young adult males were euthanized in accordance with approved protocols. Liver tissue was collected into a conical tube and snap-frozen in liquid nitrogen. Frozen livers were shipped to Dovetail Genomics for DNA extraction and sequencing. One liver sample was used for Dovetail PacBio sequencing, and one liver was used for Omni-C library preparation and sequencing.

Six individual samples were submitted for RNA sequencing - brain and liver tissues from a young adult male and female, as well as two embryos – one of each sex. All tissues were snap-frozen in liquid nitrogen.

### Egg collection and incubation

Veiled chameleon eggs were collected at oviposition at the Reptiles and Aquatics Facility at Stowers Institute for Medical Research. The eggs were incubated in 48oz deli cups on moistened medium vermiculite with 1:1 ratio water to vermiculite. Ambient humidity was at 95% in each incubator, and 28⁰C temperature unless otherwise noted.

### Chameleon embryonic fibroblasts

Chameleon embryos were collected at stages 26-35 of development (∼75-120 days post oviposition). Embryo sex was determined phenotypically based on the presence of hemipenes, and tissue was collected for additional DNA analysis using male-specific markers^23^. Each embryo was processed separately. First, we removed the embryonic head and liver, and most internal organs. The remaining body tissue was minced in a 60mm sterile dish with a scalpel in 2ml of trypsin (Gibco 25200056). The minced tissue was then incubated at 30⁰C for 10 min.

After the incubation we vigorously pipetted fetal tissue to obtain a single cell suspension. The volume was brought up to 10ml with 8mls of culture media (DMEM/F-12 (Gibco 10565-018), 10% fetal bovine serum, 15% chicken embryo extract^88^, 1X antibiotic antimycotic solution (Sigma Millipore A5955-100 ML) and 1µg/ml gentamycin solution (VWR 97062-974)) and plated in a 100mm dish. The cells were incubated at 30⁰C with 5% CO_2_.The media was changed 2 days later, and every 2 days thereafter. Cells were split as needed, as they became confluent.

### Nuclear propidium iodide staining and DNA analysis

Chameleon embryonic fibroblasts were expanded, and male and female cells from different embryos were pooled to obtain 1.5e6 to 3e6 cells for each sex. Genome size was determined using flow cytometry of propidium iodide stained nuclei, following published protocols^89^. Mouse bone marrow, human HAP1 and *Drosophila melanogaster* CME were used as control cell lines with known genome sizes to create the standard curve of cellular DNA concentration.

### Dovetail genome assembly

288.5 gigabase-pairs of PacBio CLR reads (Supplementary Table 7) were used as an input to WTDBG2 v2.5 with genome size 1.8g, minimum read length 20000, and minimum alignment length 8192. Additionally, realignment was enabled with the -R option and read type was set with the option -x sq. Dovetail Genomics did not perform contig polishing at the time of assembly.

Blast results of the WTDBG2^90^ output assembly (asm.cns.fa) against the nt database were used as input for blobtools v1.1.1^91^ and scaffolds identified as possible contamination were removed from the assembly (filtered.asm.cns.fa). Finally, purge_dups v1.2.3^92^ was used to remove haplotigs and contig overlaps (purged.fa).

### Dovetail Omni-C library preparation and sequencing

For each Dovetail Omni-C library, chromatin was fixed in place with formaldehyde in the nucleus and then extracted. Fixed chromatin was digested with DNAse I, chromatin ends were repaired and ligated to a biotinylated bridge adapter followed by proximity ligation of adapter containing ends. After proximity ligation, crosslinks were reversed, and the DNA purified.

Purified DNA was treated to remove biotin that was not internal to ligated fragments. Sequencing libraries were generated using NEBNext Ultra enzymes and Illumina-compatible adapters. Biotin-containing fragments were isolated using streptavidin beads before PCR enrichment of each library. The library was sequenced on an Illumina HiSeqX platform to produce an approximately 30x sequence coverage. Then HiRise^33^ used MQ>50 reads for scaffolding.

### Scaffolding the assembly with HiRise

The input de novo assembly and Dovetail OmniC library reads were used as input data for HiRise, a software pipeline designed specifically for using proximity ligation data to scaffold genome assemblies^33^. Dovetail OmniC library sequences were aligned to the draft input assembly using bwa^93^ (https://github.com/lh3/bwa). The separations of Dovetail OmniC read pairs mapped within draft scaffolds were analyzed by HiRise to produce a likelihood model for genomic distance between read pairs, and the model was used to identify and break putative misjoins, to score prospective joins, and make joins above a threshold. The assembly was reviewed and manually curated to correct any misjoins and make any missed joins.

### Gene model

To generate gene models, we ran Helixer v.0.3.4^39,40^ using only the genome assembly. We further supplemented Helixer-generated models with previously published IsoSeq transcriptome^12^ using agat_sp_complement_annotations.pl (v0.9.1)^94^. The IsoSeq transcriptome GFF was generated using minimap2(v.2.26)^95^, Trinity(v.2.15.1)^96^ and TransDecoder (v.5.5.0)^97^. Additionally, some genes that were not identified computationally, were identified and annotated manually using Apollo^98^.

Orthologs were identified using standalone Orthologous Matrix (OMA)(v 2.6.0 and Jul_2023 db release)^99^. Gene names were assigned using OMA orthologs, with priority given to human. If no ortholog was identified, the UNIPROT (v 2024_01) best blast hit (blastplus v2.13.0) was used.

To access the completeness of the gene model predictions and gene annotation we ran BUSCO v5.4.7^41^ on the genome and the translated gene model proteins using default parameters and the following lineages (odb10): Metazoa, vertebrate, Tetrapoda and Sauropsida.

### Chromosome identification

Scaffolds were assigned to chromosomal identities using Chromosome-specific FACS-sorted DNA sequencing data from Tishakova et al (2022) (SRA PRJNA832590)^24^. Reads from each chromosome were trimmed using TrimGalore (v0.6.6)^100^ and then aligned to our scaffolded assembly using bwa mem (v2.2.1). After alignment, our reference assembly was broken up into 100kb bins and the number of reads aligning to each of the bins was calculated using bedtools coverage (v2.30.0)^101^. After visualization, we were able to assign chromosomal identities to our scaffolds based on which scaffold the reads from each chromosome sample predominantly aligned to.

### Repeat identification and masking

After assembling the new chameleon genome, RepeatModeler (v2.0.4) was used to annotate repeat elements in the assembly. RepeatMasker (v4.1.2)^102^ was then used to softmask these repeats. After the initial masking, RepeatProteinMask from RepeatModeler was used with bedtools maskfasta to mask repeat proteins found across the genome.

One specific repeat element of interest, the interstitial telomeric repeat (TTAGGG)_n_, was searched for across the genome using seqkit (v.2.3.1)^103^. The specific sequence searched for was (TTAGGG)_4_ and one mismatch was allowed. The number of repeats found in each 100kb of the assembly was calculated using bedtools coverage.

### Genome metrics

After the genome was adjusted to separate the microchromosomes, the values for the following metrics were recalculated using QUAST (v.5.2.0)^104^: Lx, Nx, Total genome length, and Number of Contigs.

The genome assembly was filtered to only focus on the 12 primary chromosomes (6 macro; 6 micro) and then each chromosome was broken up into 100kb bins for calculating a variety of metrics. Gene Density was calculated by intersecting the 100kb bins with our gene models and finding the number of genes per bin. Repeat Density was calculated in the same way, but using the repeats annotated using RepeatModeler instead of genes. Gaps were defined as any genomic region with 10 or more consecutive Ns. The number for each of these three metrics was calculated using bedtools coverage across the 100kb bins. The final metric, - percent GC, was calculated using bedtools nuc for the same 100kb bin regions. After calculating these metrics, they were plotted on a genome-wide scale using ggplot2 (v3.4.3; R v4.2.3)^105^.

To further explore the GC content peaks, jellyfish (v2.2.7)^106^ was used to generate overrepresented kmers (both 6mers and 21mers were used) in the genomic regions surrounding (and including) the GC peaks. Seqkit was then used to scan the genome and find all occurrences of these kmers. Bedtools coverage was once again used to calculate the number of occurrences in each genomic bin. The kmers were then ranked by calculating the correlation between the number of occurrences of that specific kmer and the GC content of each bin.

### Synteny analysis

Synteny analysis was run between the finalized veiled chameleon, the original (pre-split) veiled chameleon, chicken (GCF_000002315.5), brown anole (fetched from https://dataverse.harvard.edu/dataset.xhtml?persistentId=doi:10.7910/DVN/TTKBFU), panther chameleon (fetched from https://doi.org/10.57760/sciencedb.08450), and dwarf chameleons *Bradypodion pumilum* (GCA_035047305.1) and *Bradypodion ventrale* (GCA_035047345.1) assemblies using Progressive Cactus (v2.4.0)^107^. The required phylogenetic tree input for cactus was generated using Mashtree (v.1.2.0)^108^. After finding the pairwise syntenic blocks between the finalized Veiled Chameleon assemblies and the other genomes, syntenyPlotteR (v1.0.0)^109^ was used to generate both the In-Silico Chromosomal Painting and Synteny Sankey plots.

### Sex chromosome analysis

The determination of the male-specific sex determination region was primarily done using two parallel approaches focused on previously published data^23^. The first involved aligning male-specific markers using blastn (v.2.12.0)^110^. The best hit for each marker was found by ranking the hits based on bit score and keeping the highest score. The second method involved processing the RAD-seq data to derive the sex determination region. The raw RAD-seq data was fetched from SRA Accession PRJNA429428. Reads were then aligned using minimap2 (v2.26)^95^ and sorted with samtools (v1.18)^111^. Stacks (v.2.55)^112^ was used to call SNPs and generate summary statistics comparing nucleotide diversity between the male and female samples. The Smoothed F_ST_ value was averaged across each 100kb bin and plotted across the genome.

Additionally, we compared the overlap between the non-recombining regions of the sex chromosomes in *C. calyptratus* and *E. heatwolei* by blasting Y-linked transcripts from *E. heatwolei*. Y-linked transcripts were aligned to our assembly using blastp (v.2.12.0), keeping the best hit based on the highest bit score. The location of these alignments was then compared to our putative sex determination region.

### Environment and sex determination

3 clutches of eggs from different females, laid in May 2018 were used for the experiment. The eggs were incubated in 48oz deli cups according to standard protocols. Each container was labeled with numbers on the side as well as on the lid so each egg could be identified individually. Eggs were first incubated at 28⁰C, and then subdivided into the 4 temperature groups (24⁰C – 49 eggs, 26⁰C – 47 eggs, 28⁰C – 48 eggs and 30⁰C – 48 eggs) at 70days post oviposition – end of gastrulation, prior to known sex determination processes. The eggs were incubated for 90 days until 160 days post oviposition. The eggs were weighed at the start and end of the project prior to their dissection at 160 days post oviposition. Only 2/192 eggs were non-viable.

The embryos were dissected out of the eggs at 160 days post oviposition, and a piece of tissue preserved for DNA genotyping. The phenotypic sex was determined independently by two investigators based on the presence of heel spur or hemipenes. The animals that received different sex assignments by the two investigators were re-assessed until the consensus was reached for all animals. The DNA genotyping, using previously published sex-specific PCR primers^23^, was carried out by an investigator, blinded to the results of the phenotypic assessment. Finally, the results of phenotypic and genetic sex determination were aligned to reveal 5 genotypic females, which were assessed as males phenotypically.

### Statistical analysis

We used Prism-GraphPad to carry out statistical analyses where appropriate.

## AKNOWLEDGEMENTS

The authors thank members of the Trainor lab and the Computational Biology center for their insights and discussion throughout the course of this project, especially Madelaine Gogol and Eric Ross for sharing their experience and expertise in genome analysis and annotation. Michay Diez was exceptionally helpful in our discussion about rDNA regions in the genome. We greatly appreciate Alex Muensch, David Jewell, Kristy Winter, Christina Piraquive, Diana Baumann, Elizabeth Evans and the Reptile Facility for their care, husbandry, and maintenance of our veiled chameleon colony. This work was supported by the Stowers Institute for Medical Research (P.A.T) and a K99 (HD114881) from the National Institute for Child Health and Human Development (N.A.S).

---

**Supplementary Table 1 | Cross-reference of the naming conventions used in this study for chromosomes and scaffolds, as well as other studies.** For studies identifying gene location based on genomic fluorescent *in situ* hybridization, gene’s location is identified in the table in accordance with assembled genome data.

**Supplementary Table 2 | Analysis of 6-mer distribution across the genome, in correlation with GC content.** 10 sequences each which have positive, negative and no correlation with GC content across the genome. Example graphical representations available in Supplementary Figure S10.

**Supplementary Table 3 | BUSCO scores for genome and protein predictions.**

**Supplementary Table 4 | Locations of male-specific markers on chromosome 5.** Markers were originally identified through RAD-seq of male and female genomes^23^.

**Supplementary Table 5 | Predicted genes in the sex determination region on chromosome 5.** Genes highlighted in gray overlap the first 11 male-specific markers in the first 0.85Mb of chromosomes 5. Notes column supplies additional information regarding potential involvement of specific genes in sex determination and fertility. PMID indicates the publication referencing information in the Notes column.

**Supplementary Table 6 | Permanent gene IDs for genes mentioned in text.**

**Supplementary Table 7 | PacBio reads.**

## Notes

### Competing Interest Statement

The authors have declared no competing interest.

