## Supplementary Figures for "*Chamaeleo calyptratus* (veiled chameleon) chromosome-scale genome assembly and annotation provides insights into the evolution of reptiles and developmental mechanisms"

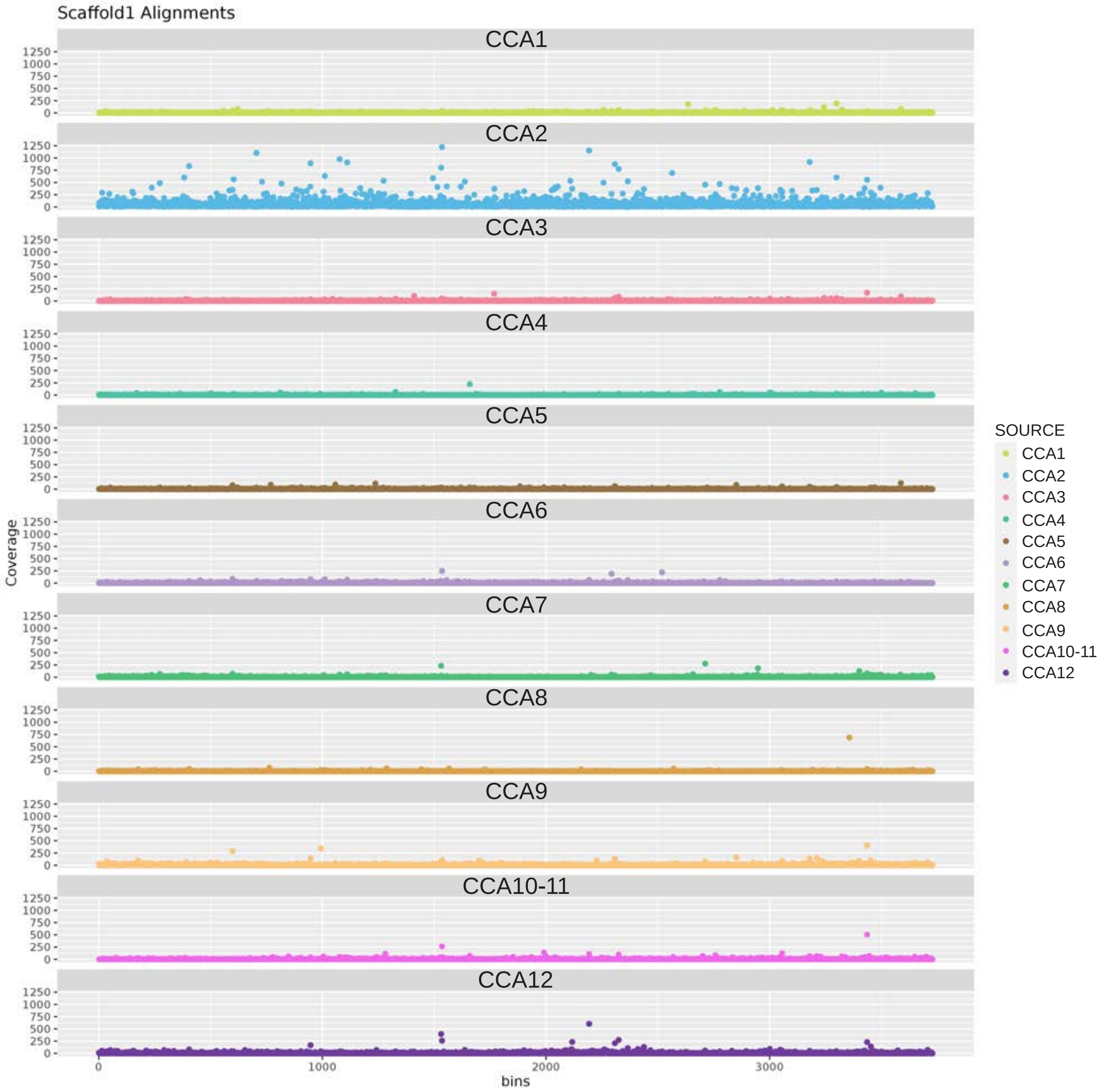

**Supplementary Figure S1 | Scaffold 1 is chromosome 2 (CCA2).** The horizontal axis represents locations on scaffold 1, binned into 100kb individual bins for mapping purposes. The vertical axis represents the number of reads. Reads from individual chromosomes were mapped to scaffold 1 (Tishakova et al., 2022). Of all chromosomal reads, most of scaffold 1 corresponds to reads from chromosome 2.

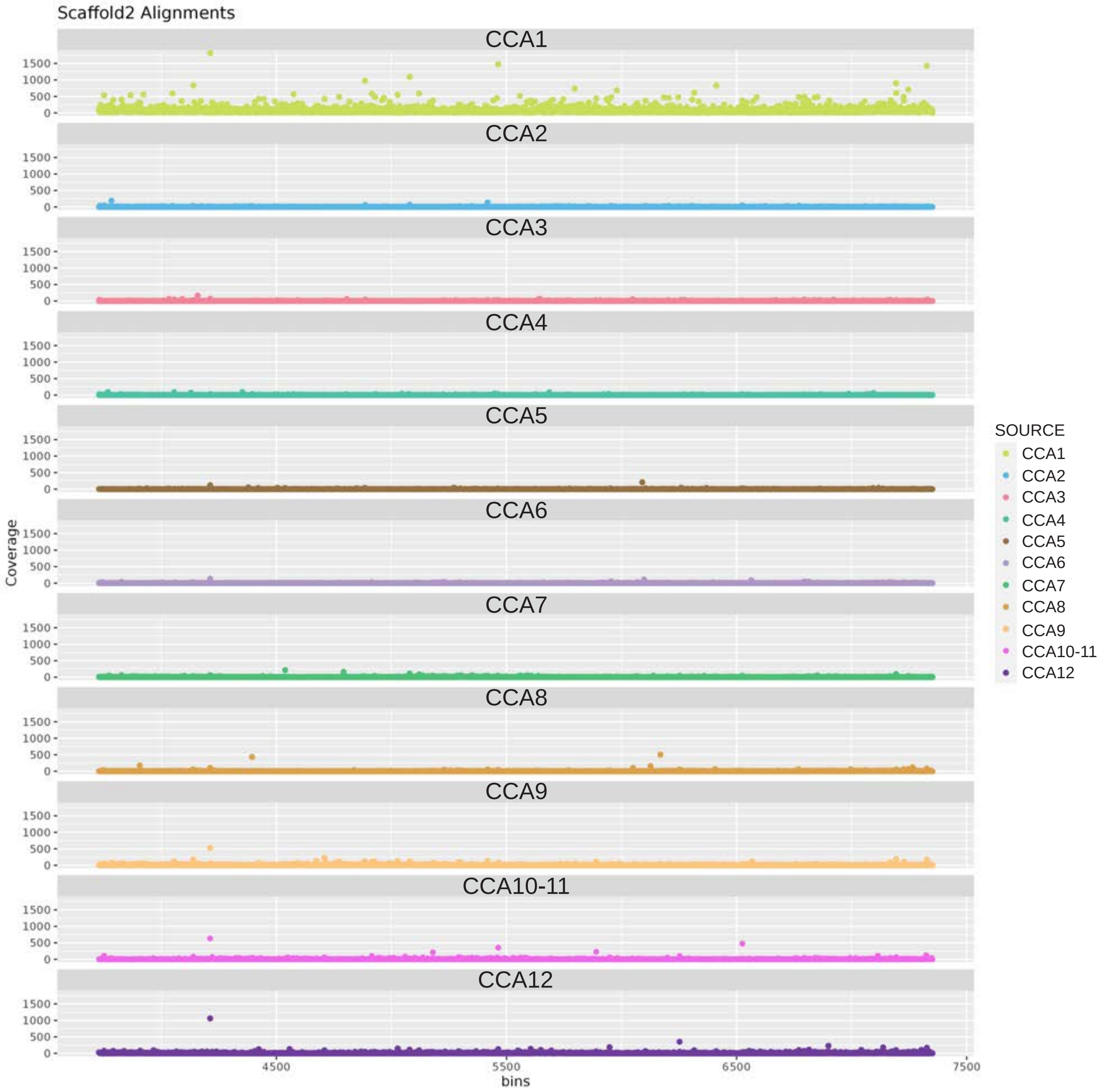

**Supplementary Figure S2 | Scaffold 2. is chromosome 1 (CCA1)** The horizontal axis represents locations on scaffold 2, binned into 100kb individual bins for mapping purposes. The vertical axis represents the number of reads. Reads from individual chromosomes were mapped to scaffold 2 (Tishakova et al., 2022). Of all chromosomal reads, most of scaffold 2 corresponds to reads from chromosome 1.

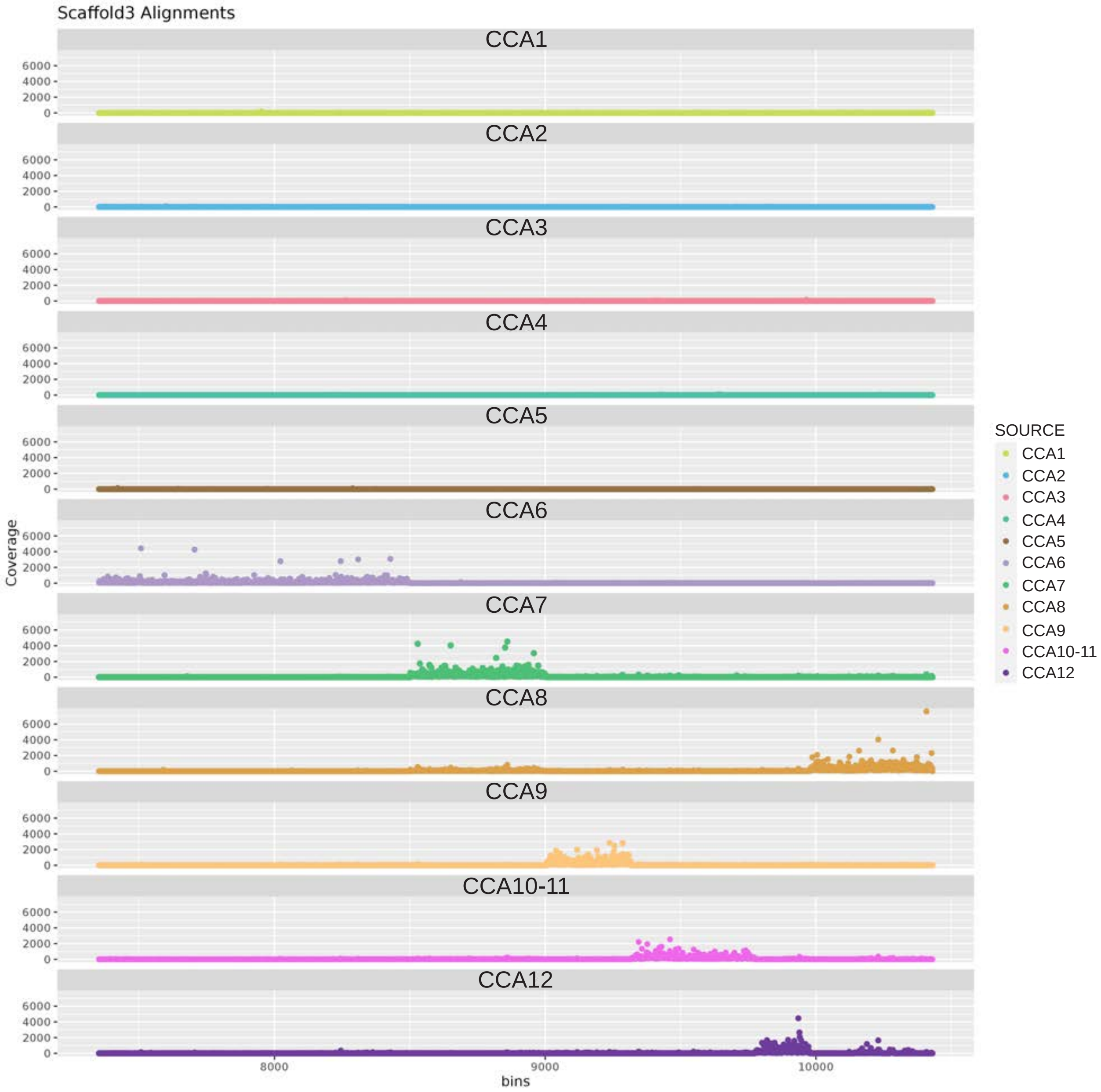

**Supplementary Figure S3 | Chromosomes 6-12 (CCA6-12) map to scaffold 3.** The horizontal axis represents locations on scaffold 3, binned into 100kb individual bins for mapping purposes. The vertical axis represents the number of reads. Reads from individual chromosomes were mapped to scaffold 3 (Tishakova et al., 2022). Reads form individual chromosomes, as well as chromosomes 10 and 11 map in defined clusters to scaffold 3, which was assembled computationally. The individual chromosomes had to be separated manually based on GC content, read coverage, synteny analysis and presence of gaps in the assembly.

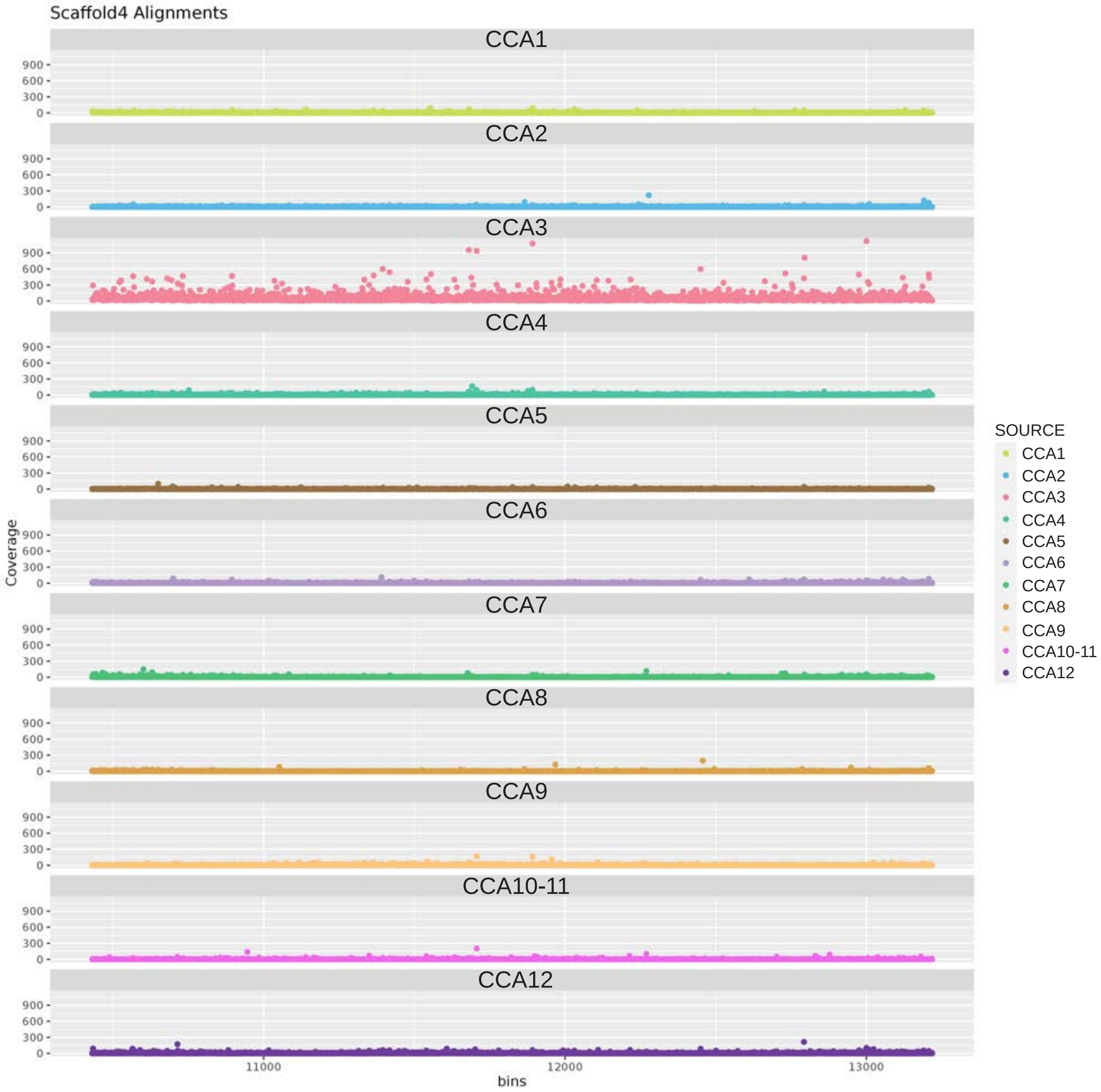

**Supplementary Figure S4 | Scaffold 4 is chromosome 3 (CCA3).** The horizontal axis represents locations on scaffold 4, binned into 100kb individual bins for mapping purposes. The vertical axis represents the number of reads. Reads from individual chromosomes were mapped to scaffold 4 (Tishakova et al., 2022). Of all chromosomal reads, most of scaffold 4 corresponds to reads from chromosome 3.

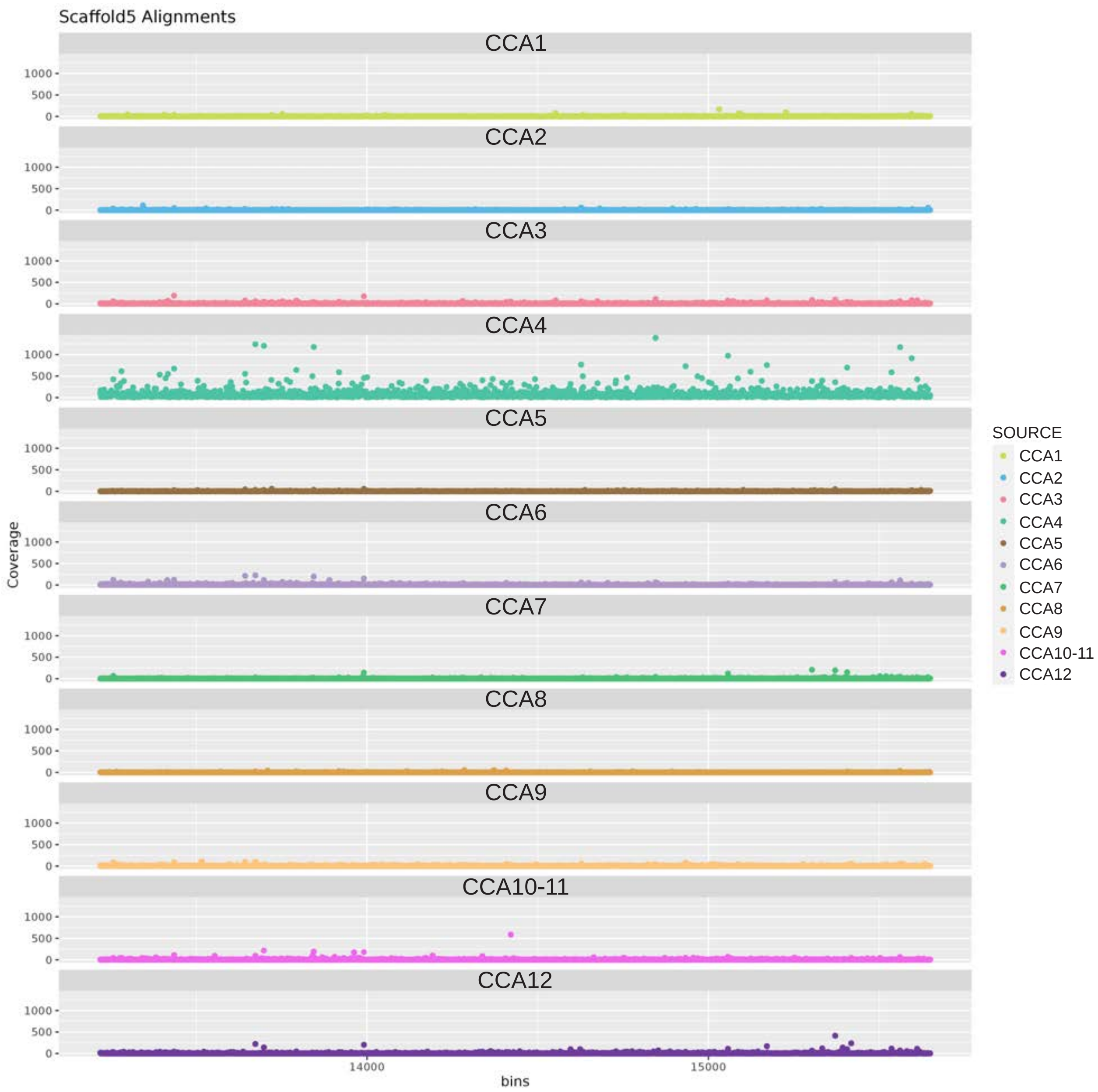

**Supplementary Figure S5 | Scaffold 5 is chromosome 4 (CCA4).** The horizontal axis represents locations on scaffold 5, binned into 100kb individual bins for mapping purposes. The vertical axis represents the number of reads. Reads from individual chromosomes were mapped to scaffold 5 (Tishakova et al., 2022). Of all chromosomal reads, most of scaffold 5 corresponds to reads from chromosome 4.

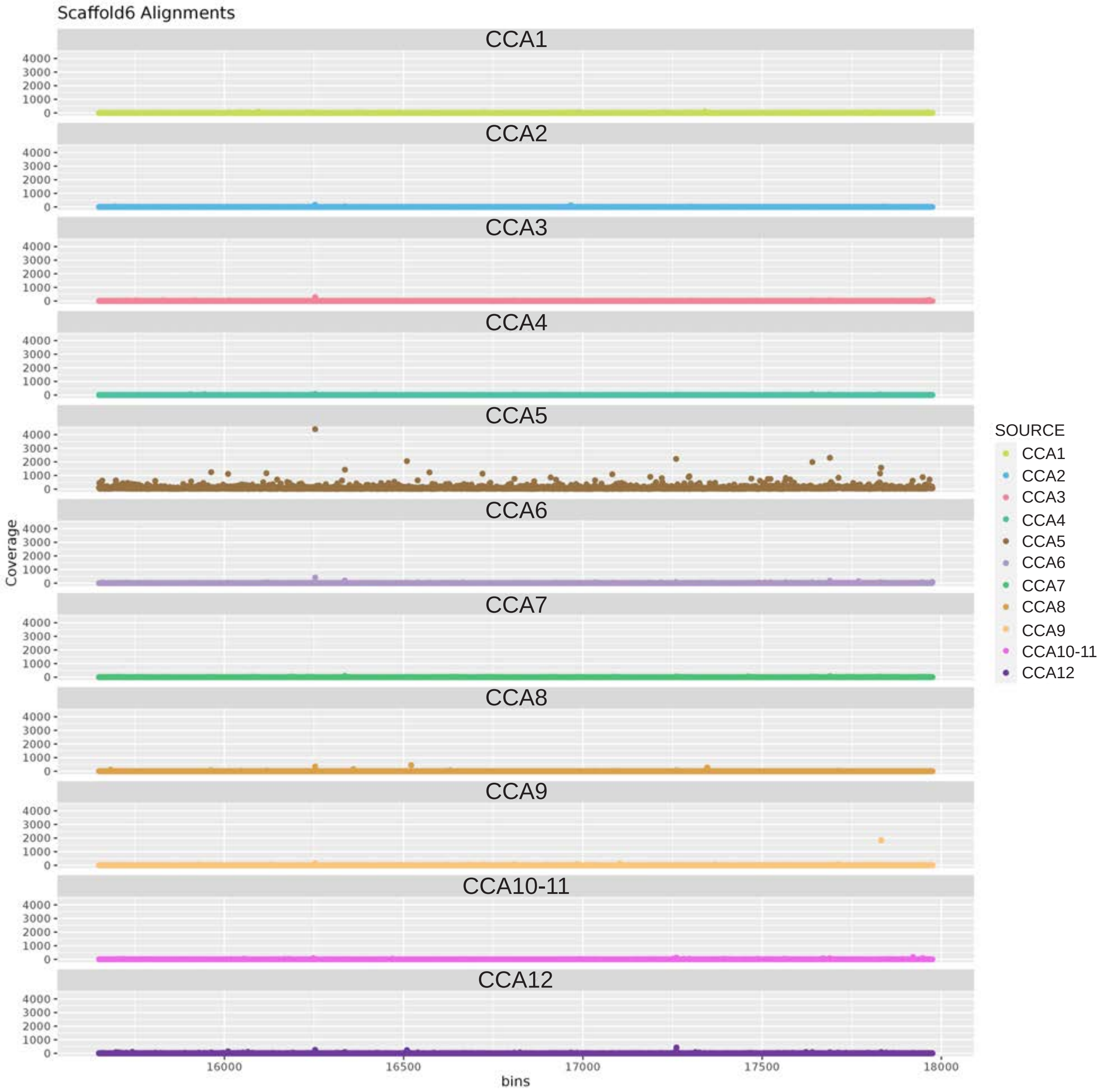

**Supplementary Figure S6 | Scaffold 6 is chromosome 5 (CCA5).** The horizontal axis represents locations on scaffold 6, binned into 100kb individual bins for mapping purposes. The vertical axis represents the number of reads. Reads from individual chromosomes were mapped to scaffold 6 (Tishakova et al., 2022). Of all chromosomal reads, most of scaffold 6 corresponds to reads from chromosome 5.

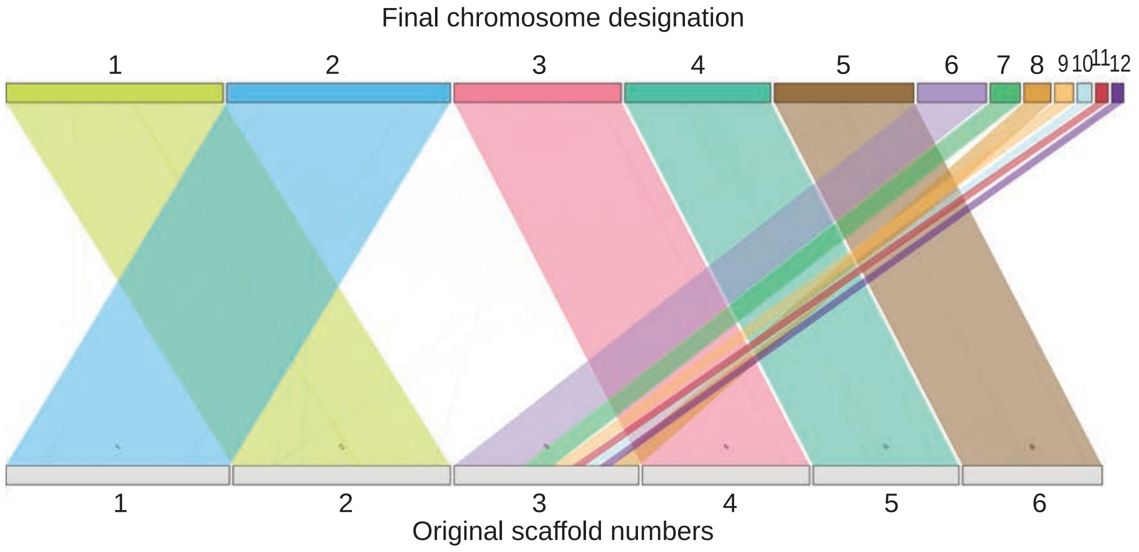

**Supplementary Figure S7 | Relationship between final chromosome designations and original scaffolds.**

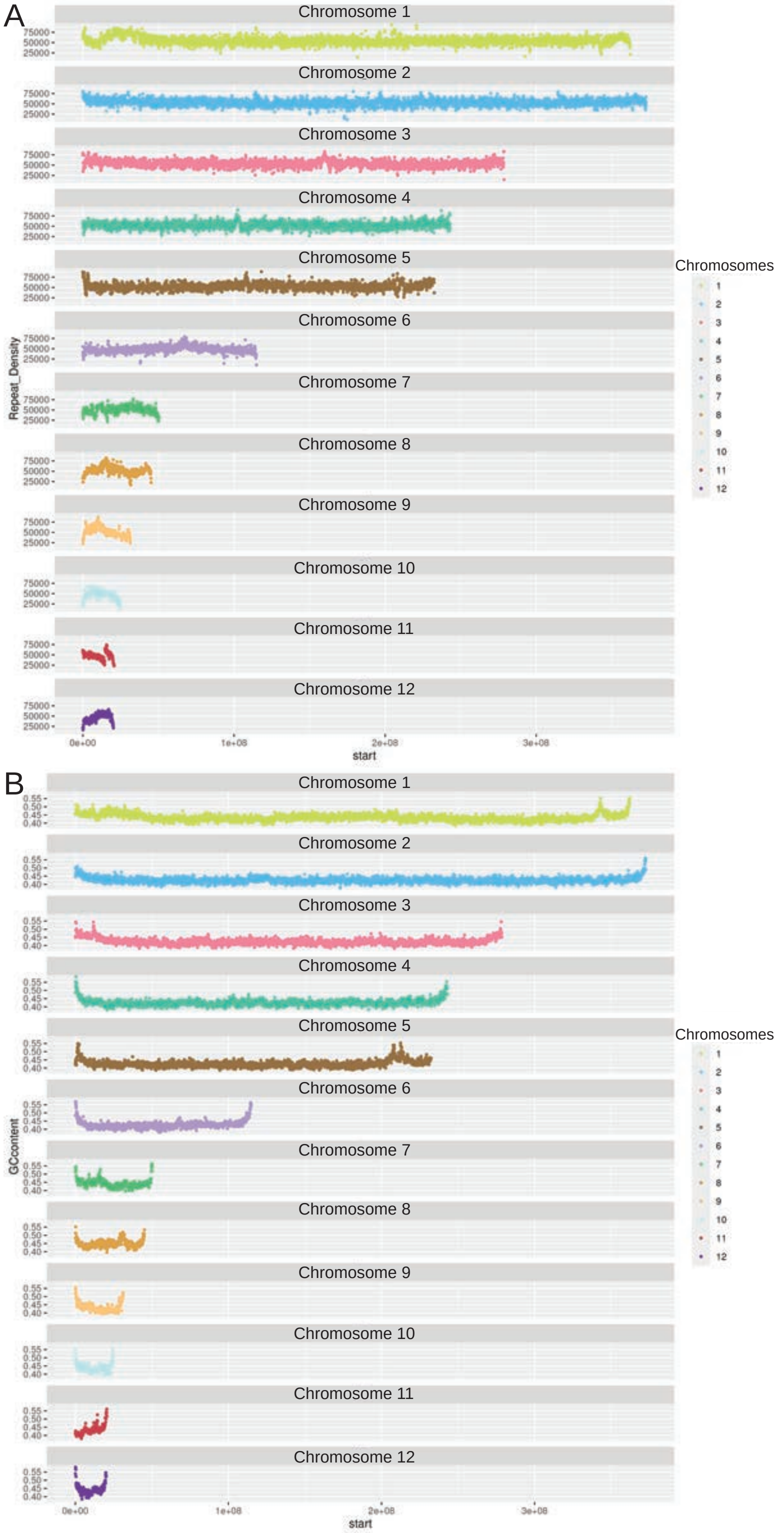

**Supplementary Figure S8 | Repeat and GC content on individual chromosomes.** (A) Repeat content on individual chromosomes. (B) GC content on individual chromosomes. The horizontal axis represents chromosomal locations, binned into 100kb individual bins for mapping purposes. The vertical axis represents the proportion of repeats or GC content in each individual bin.

Interstitial Telomeric Motif $(TTAGGG)_n$

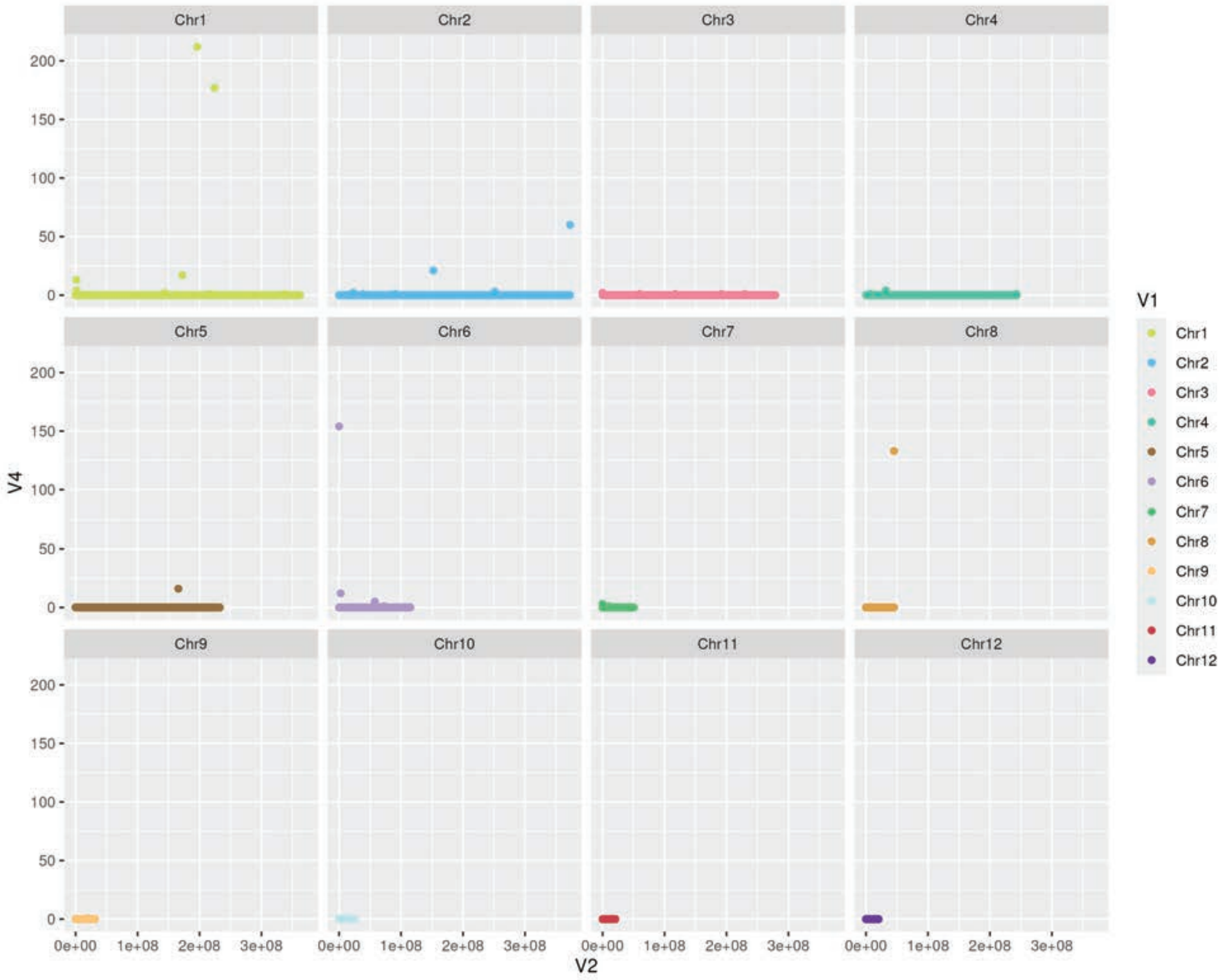

**Supplementary Figure S9 | Distribution of Interstitial Telomeric Motif (TTAGGG)n.** The highest number of repeats was observed in the pericentromeric region of chromosome 1, which is consistent with findings by Rovatsos et al (2015). The horizontal axis represents chromosomal locations, subdivided into 100kb bins for mapping purposes. The vertical axis represents the number of repeats per bin.

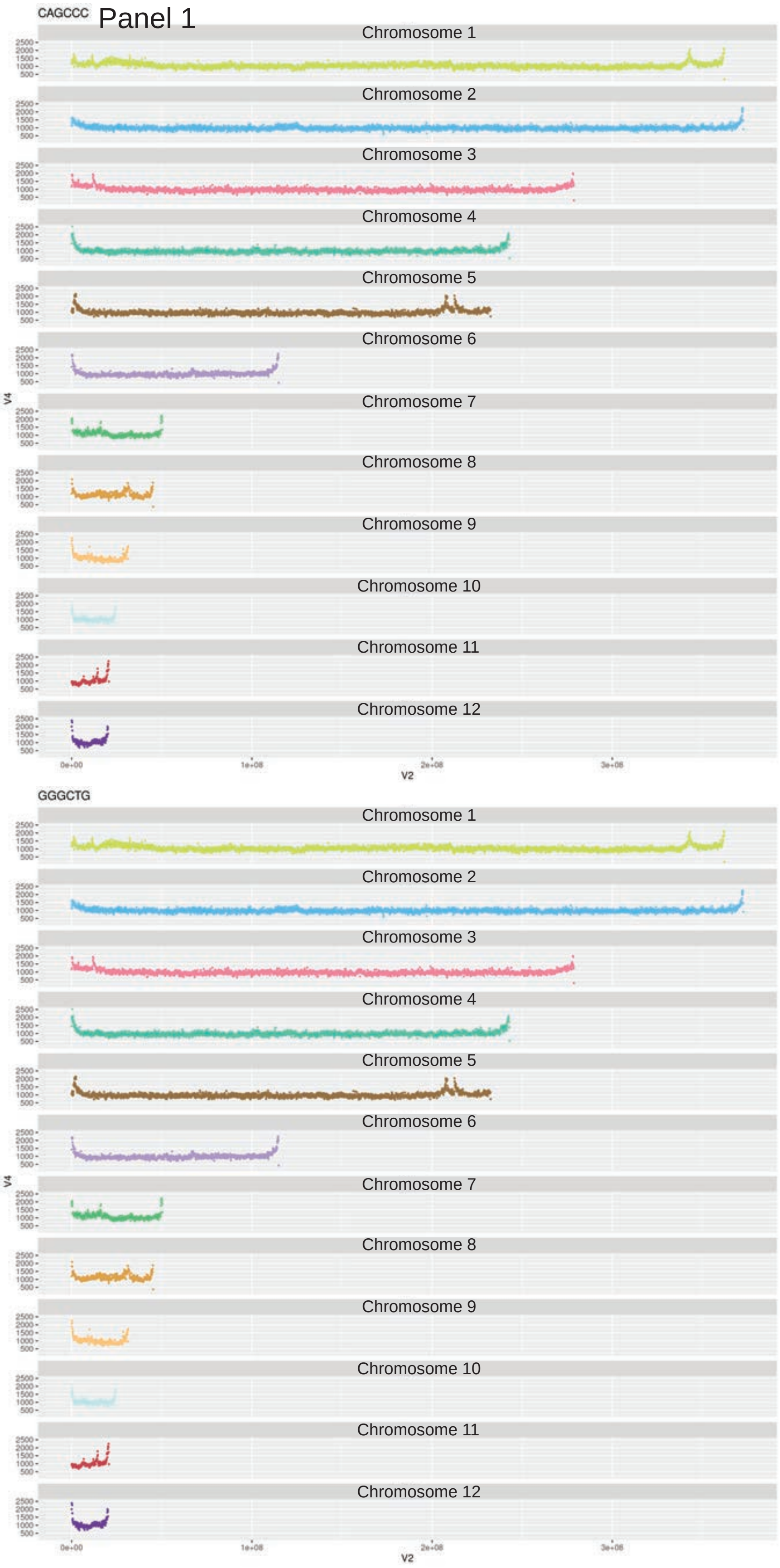

**Supplementary Figure S10 | Distribution of 6-mers that correlate positively, negatively or have no correlation with GC content in the genome.** (Panel 1) Example of positive correlation with GC content: CAGCCC/GGGCTG (correlation coefficient = 0.948018); (Panel 2) Example of negative correlation with GC content: ATTTCA/TGAAAT (correlation coefficient = -0.95159); (Panel 3) Example of no correlation with GC content: ACGACA/TGTCGT (correlation coefficient = -0.00346). The horizontal axis represents chromosomal locations, subdivided into 100kb bins for mapping purposes. The vertical axis represents the number of repeats per bin.

Panel 2

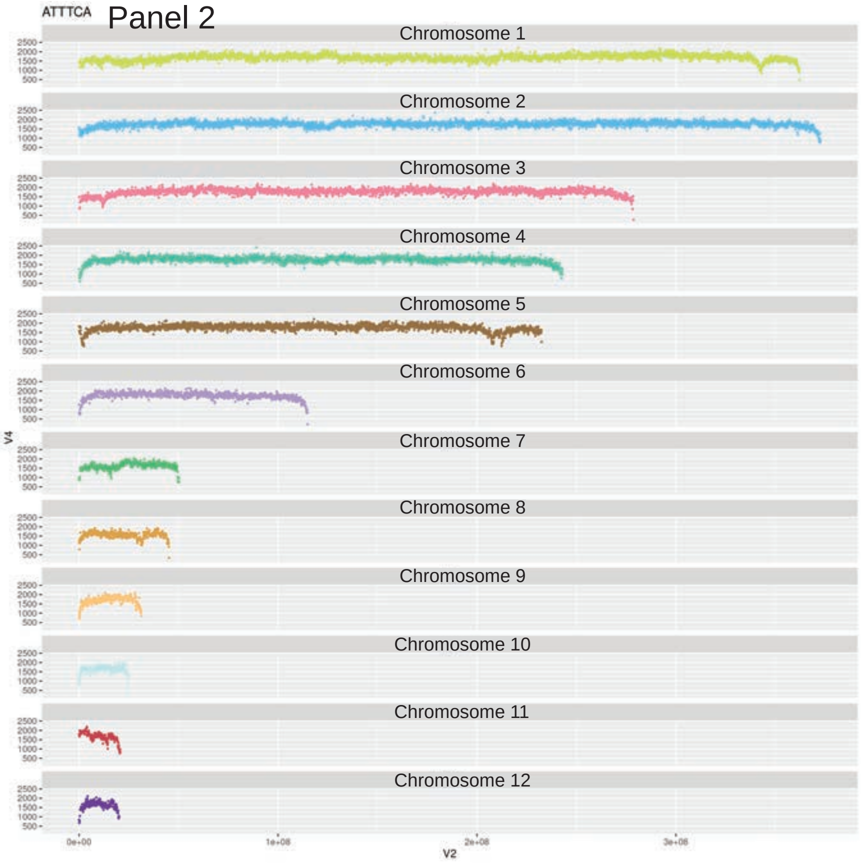
ATTTCA
Chromosome 1
Chromosome 2
Chromosome 3
Chromosome 4
Chromosome 5
Chromosome 6
Chromosome 7
Chromosome 8
Chromosome 9
Chromosome 10
Chromosome 11
Chromosome 12
V4
V2

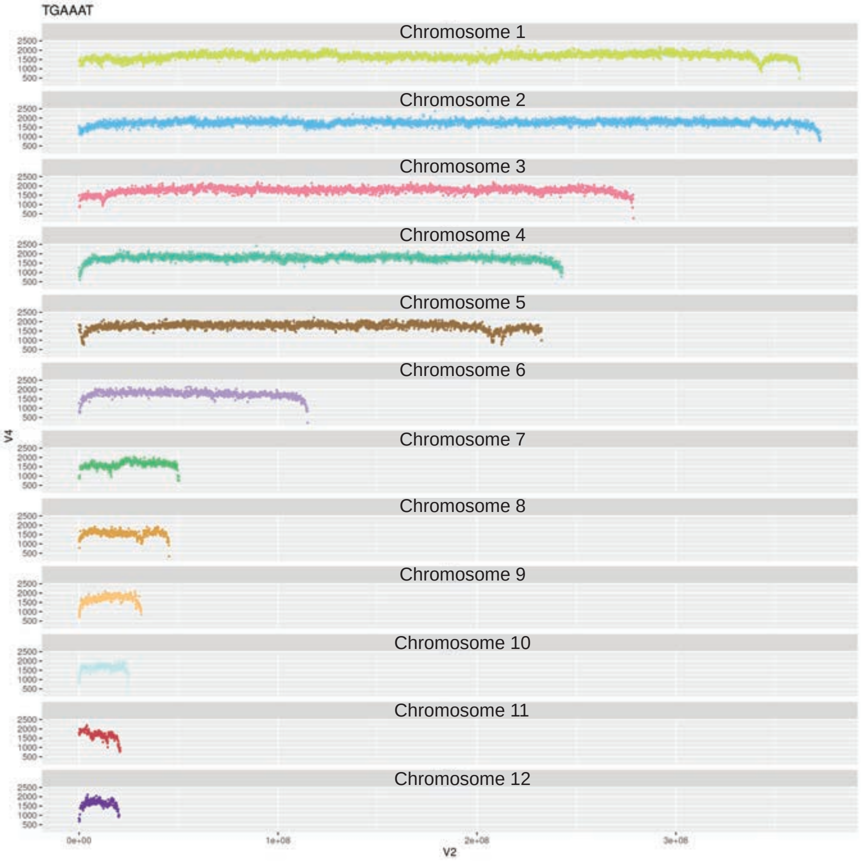
TGAAAT
Chromosome 1
Chromosome 2
Chromosome 3
Chromosome 4
Chromosome 5
Chromosome 6
Chromosome 7
Chromosome 8
Chromosome 9
Chromosome 10
Chromosome 11
Chromosome 12
V4
V2

Panel 3

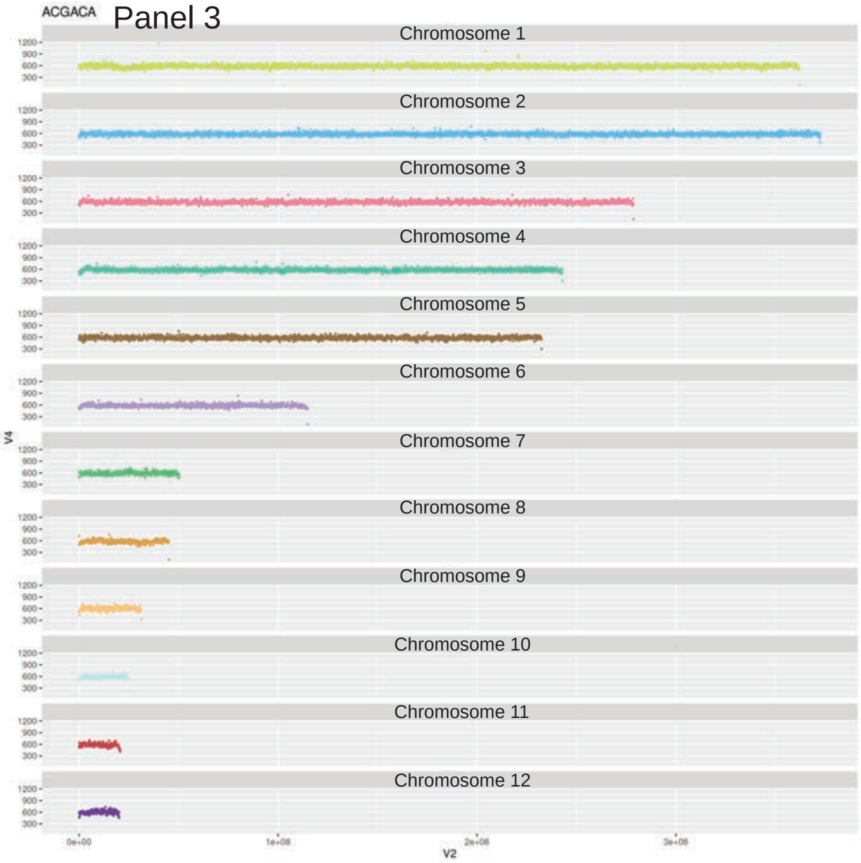
ACGACA
Chromosome 1
Chromosome 2
Chromosome 3
Chromosome 4
Chromosome 5
Chromosome 6
Chromosome 7
Chromosome 8
Chromosome 9
Chromosome 10
Chromosome 11
Chromosome 12
1200
900
600
300
V4
0e+00
1e+08
2e+08
3e+08
V2

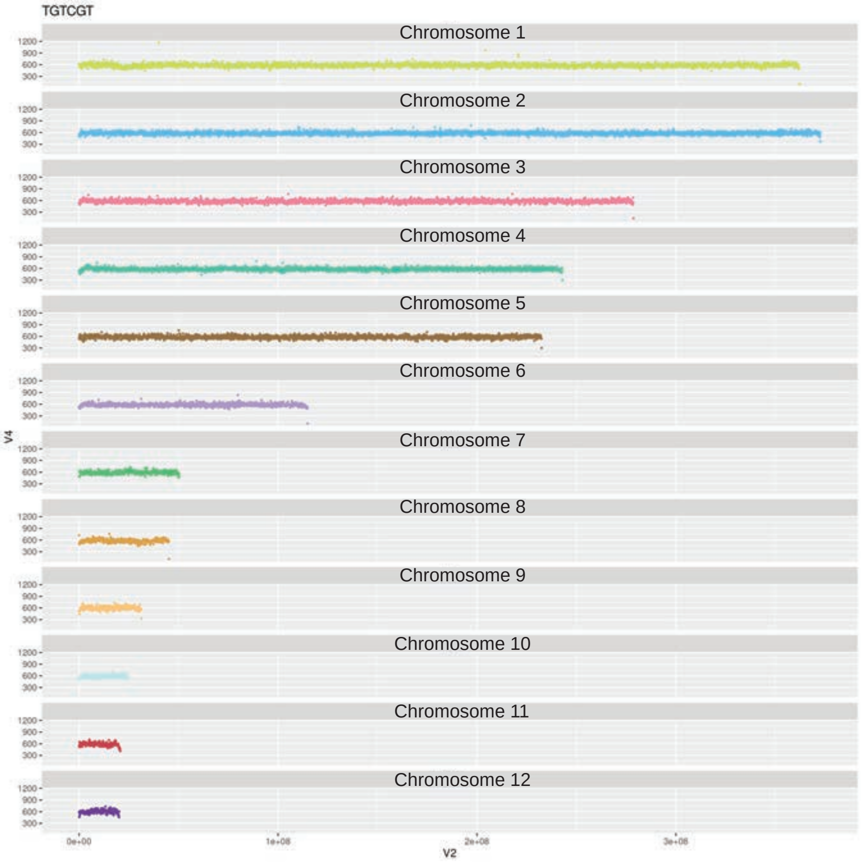
TGTCGT
Chromosome 1
Chromosome 2
Chromosome 3
Chromosome 4
Chromosome 5
Chromosome 6
Chromosome 7
Chromosome 8
Chromosome 9
Chromosome 10
Chromosome 11
Chromosome 12
1200
900
600
300
V4
0e+00
1e+08
2e+08
3e+08
V2

Panel 1

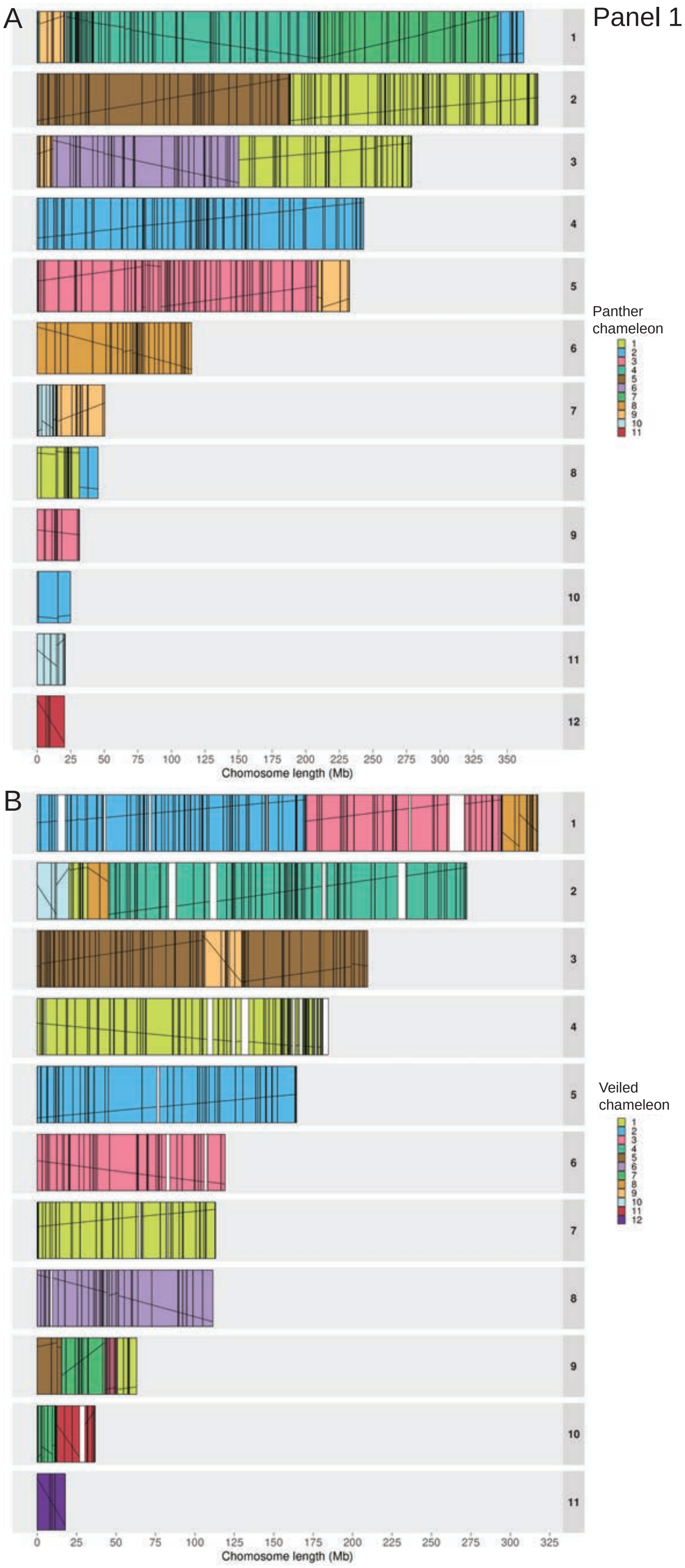

**Supplementary Figure S11 | Reciprocal synteny analysis between veiled chameleon, dwarf chameleons, panther chameleon, brown anole, and chicken.** (Panel1) Detailed synteny between veiled chameleon and panther chameleon; (Panel 2) detailed synteny between veiled chameleon and dwarf chameleon *Bradypodion ventrale*; (Panel 3) detailed synteny between veiled chameleon and dwarf chameleon *Bradypodion pumilum*; (Panel 4) detailed synteny between veiled chameleon and brown anole; (Panel 5) detailed synteny between veiled chameleon and chicken. (A) Chromosome painting, according to syntenic blocks with another species (panther chameleon, dwarf chameleons, brown anole, chicken). (B) Chromosome painting, according to syntenic blocks with veiled chameleon chromosomes. Black vertical lines denote the border regions of individual syntenic blocks. Diagonal black lines denote the co- or counter-linearity between the syntenic blocks in the two species.

Panel 2

A

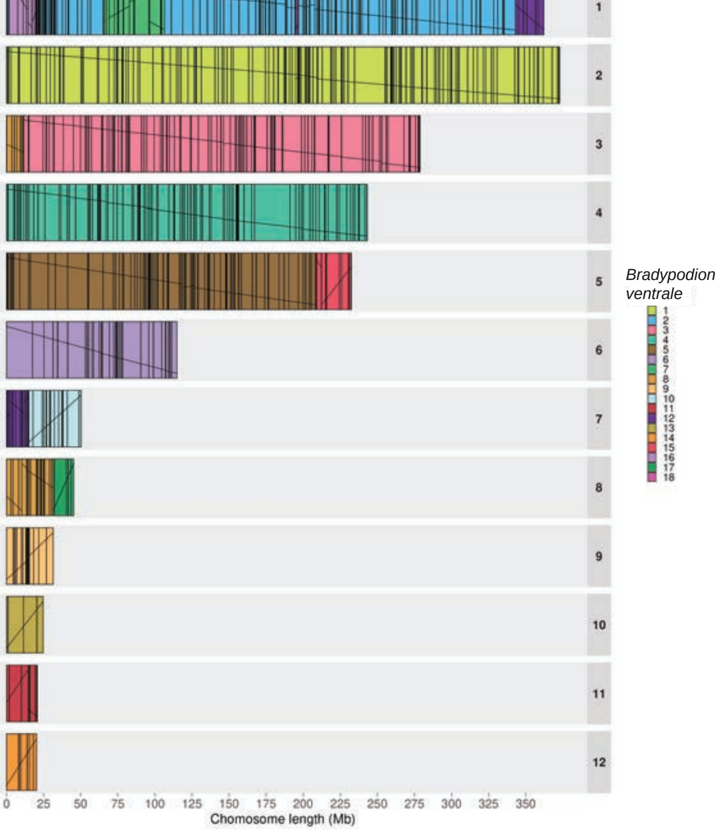
1
2
3
4
5
6
7
8
9
10
11
12
0 25 50 75 100 125 150 175 200 225 250 275 300 325 350
Chomosome length (Mb)
*Bradypodion ventrale*
1 2 3 4 5 6 7 8 9 10 11 12 13 14 15 16 17 18

B

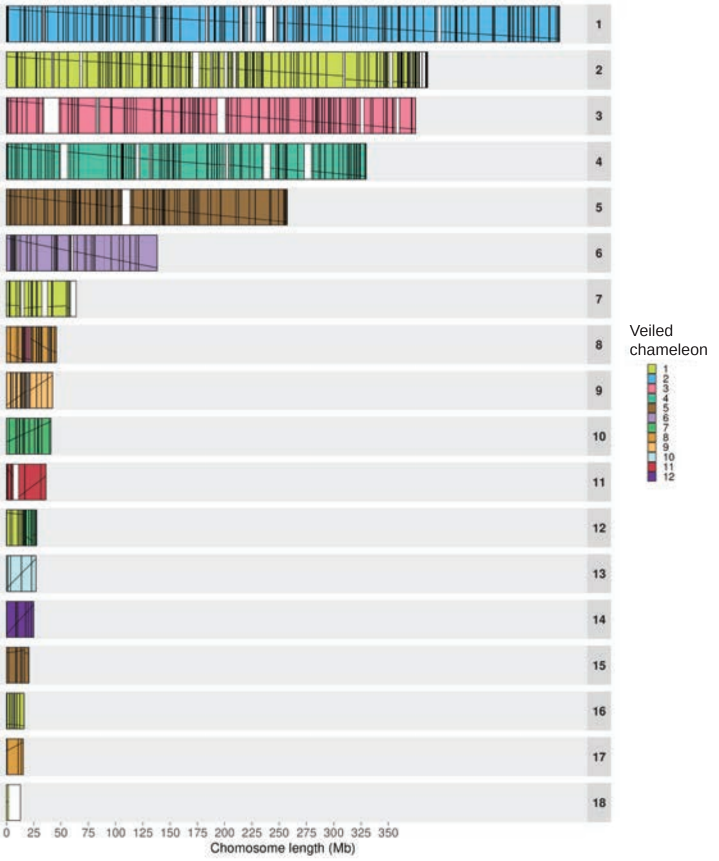
0 25 50 75 100 125 150 175 200 225 250 275 300 325 350
Chomosome length (Mb)
Veiled chameleon
1 2 3 4 5 6 7 8 9 10 11 12

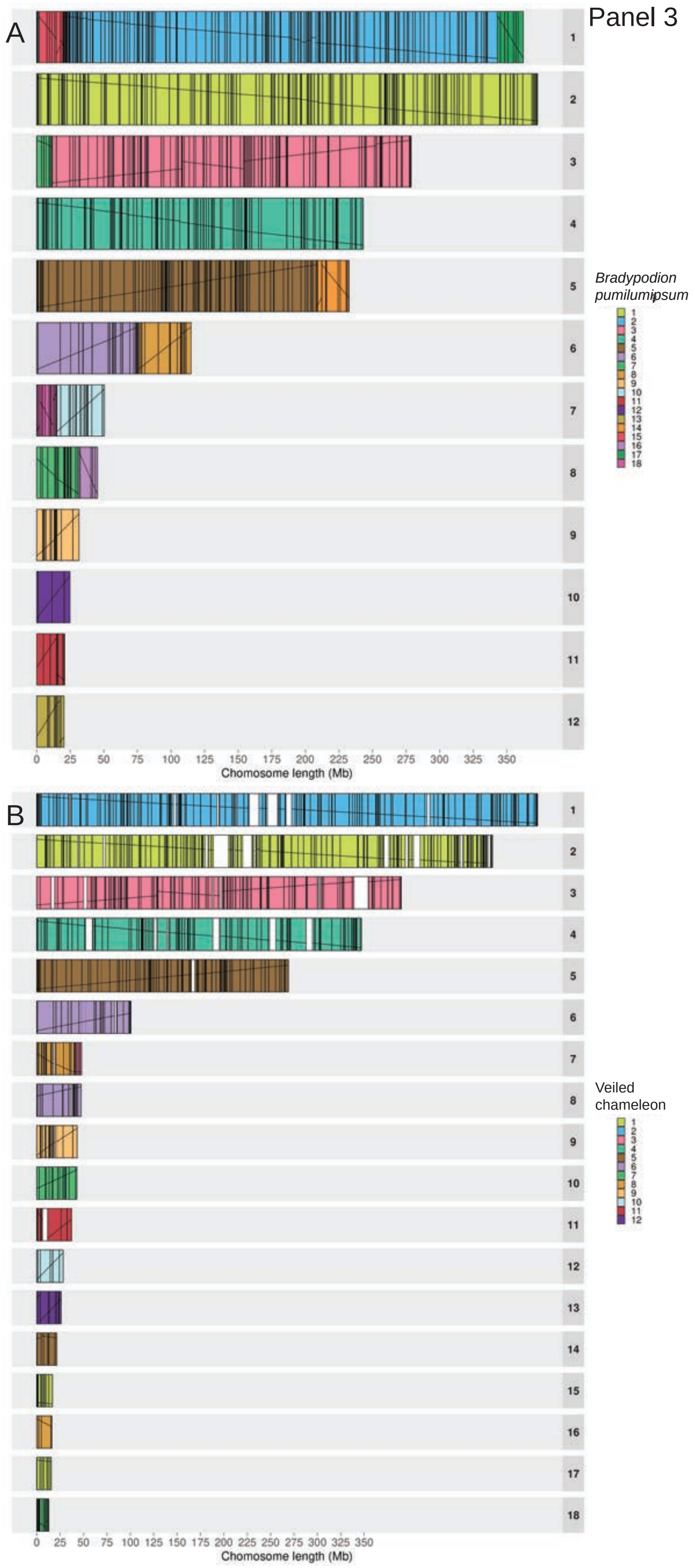
Panel 3
A
1
2
3
4
5
6
7
8
9
10
11
12
0 25 50 75 100 125 150 175 200 225 250 275 300 325 350
Chomosome length (Mb)
*Bradypodion pumilumipsum*
1 2 3 4 5 6 7 8 9 10 11 12 13 14 15 16 17 18
B
0 25 50 75 100 125 150 175 200 225 250 275 300 325 350
Chomosome length (Mb)
Veiled chameleon
1 2 3 4 5 6 7 8 9 10 11 12

A

Panel 4

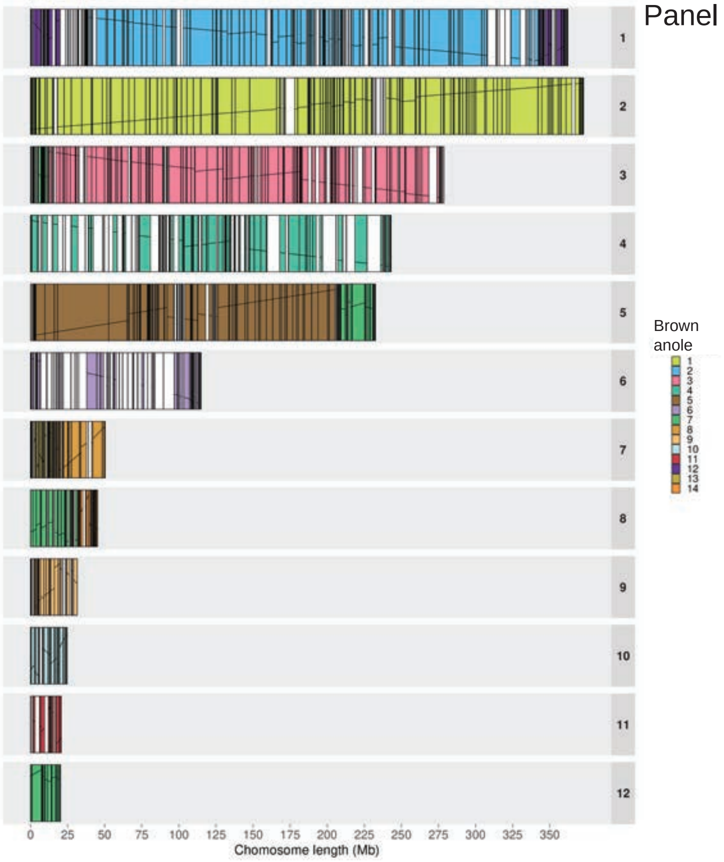
1
2
3
4
5
6
7
8
9
10
11
12
0 25 50 75 100 125 150 175 200 225 250 275 300 325 350
Chomosome length (Mb)
Brown anole
1 2 3 4 5 6 7 8 9 10 11 12 13 14

B

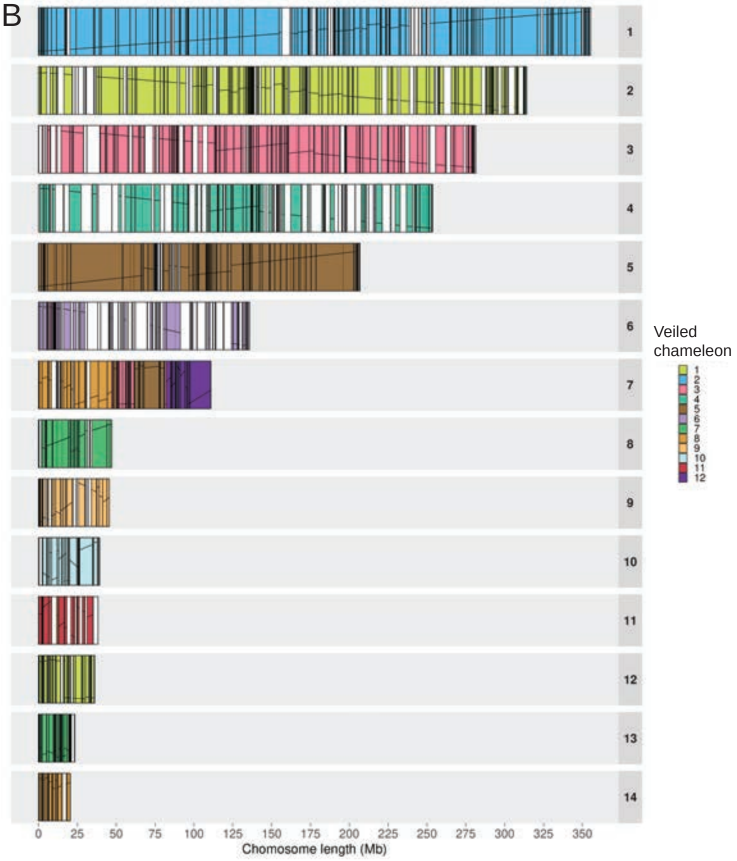
1
2
3
4
5
6
7
8
9
10
11
12
13
14
0 25 50 75 100 125 150 175 200 225 250 275 300 325 350
Chromosome length (Mb)
Veiled chameleon
1 2 3 4 5 6 7 8 9 10 11 12

Panel 5

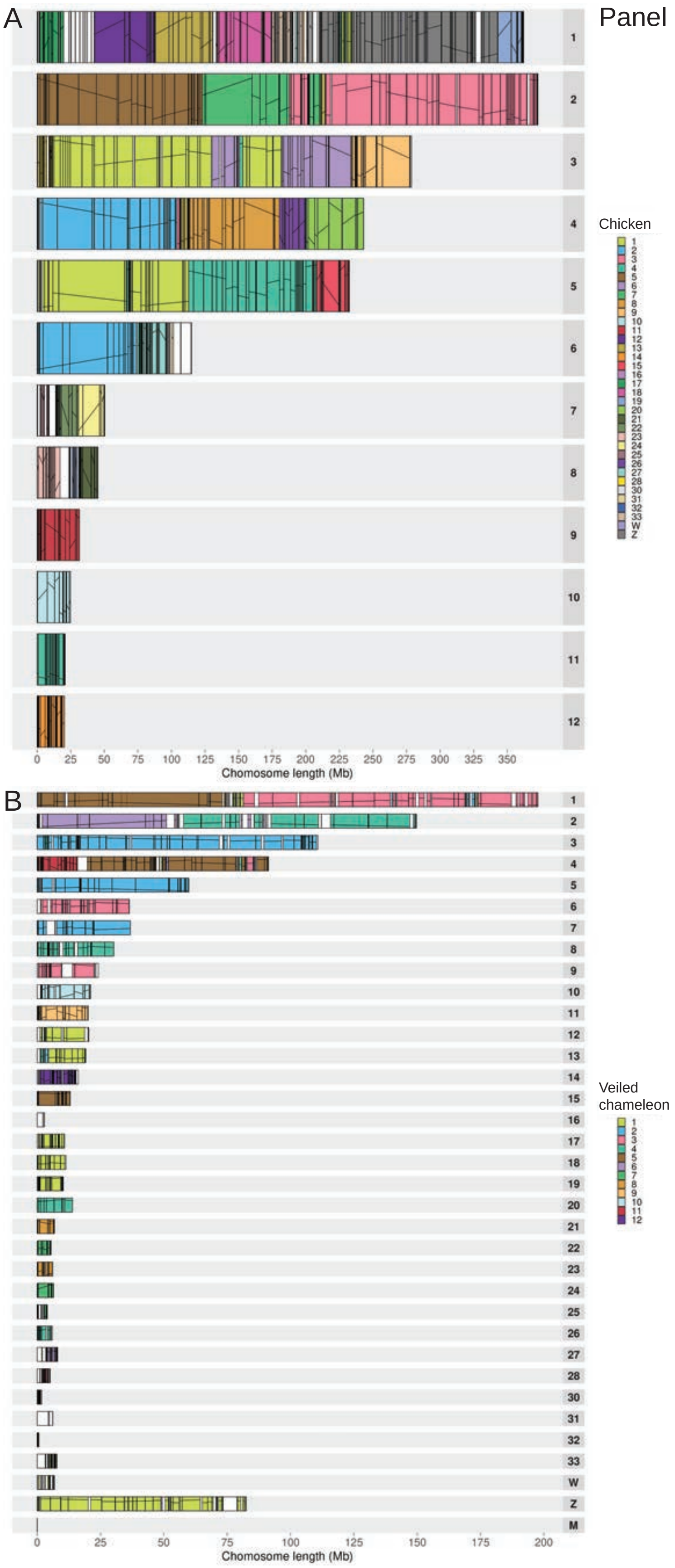
A
B
Chicken
30
31
32
33
W
Z
Chomosome length (Mb)
Veiled chameleon
M

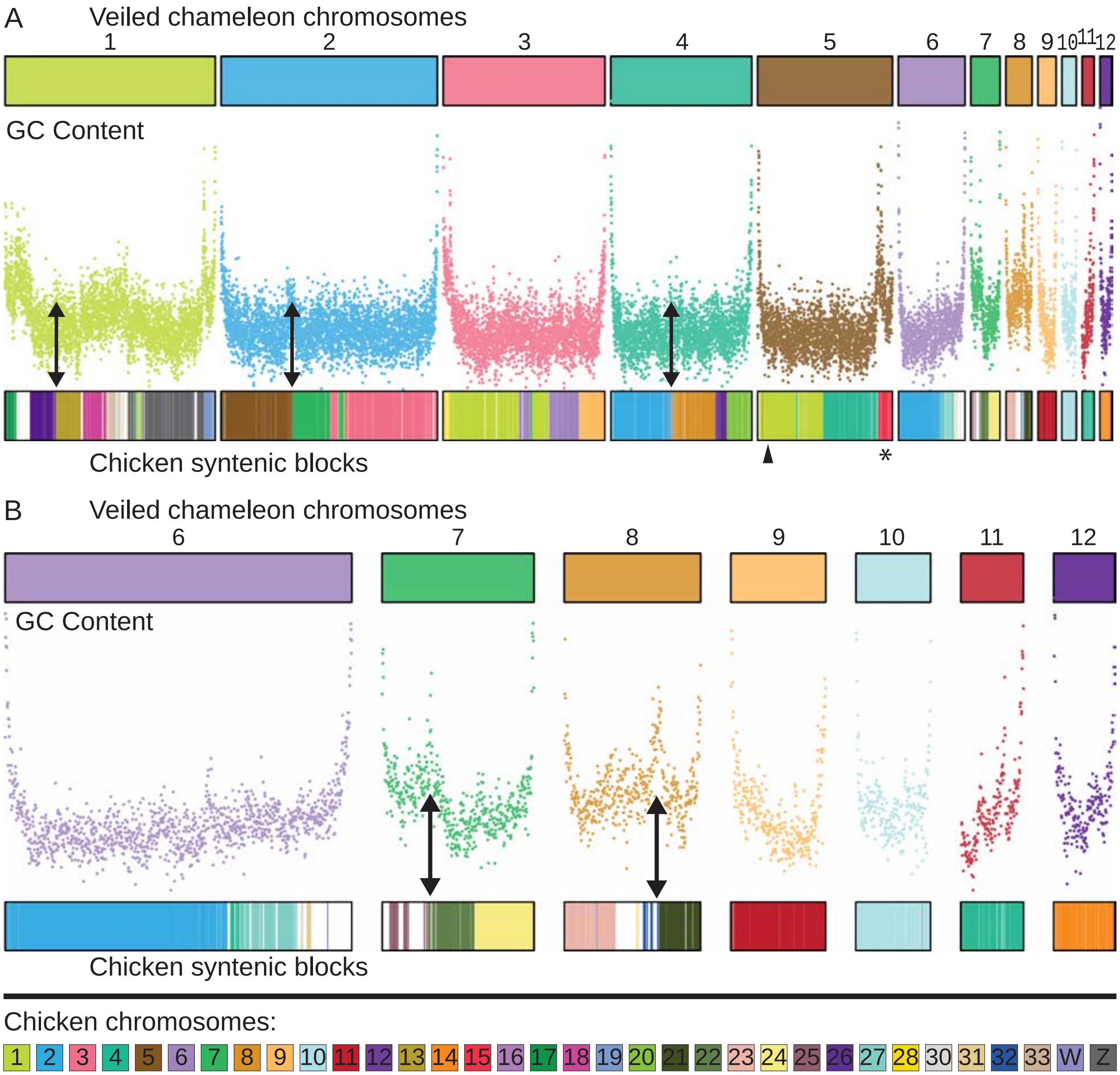

**Supplementary Figure S12 | Synteny analysis between veiled chameleon and chicken genomes.** Synteny analysis between veiled chameleon and chicken, with comparison to regional GC content. Chromosomes are painted according to synteny with chicken chromosomes. Double-headed arrows indicate several regions where fusions of chicken syntenic blocks correlate with regions of high GC content in veiled chameleon genome. (A) Chromosomal painting of the entire genome. Arrowhead denotes the location of the skink Y-linked syntenic region. Asterisk denotes the location of a genomic block, syntenic with chromosome 15 in chicken, previously identified as a conserved X-linked sex chromosome element in all pleurodont iguanas (except basilisks and their relatives; chromosome 5 in chameleon, chromosome 7 in brown anole)(Nielsen et al., 2019). (B) Chromosomal painting of chromosomes 6-12.

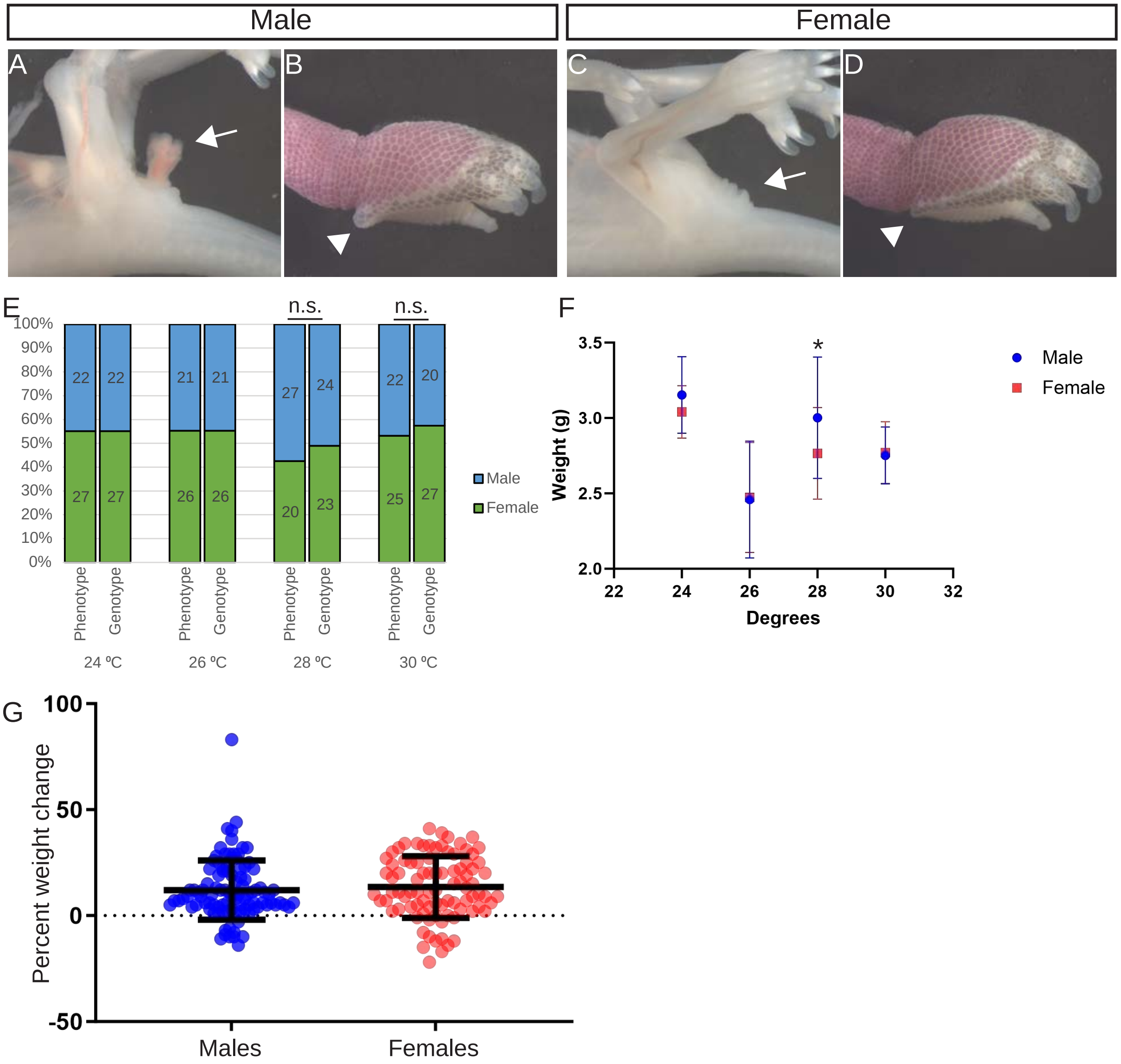

**Supplementary Figure S13 | Sex determination in veiled chameleons is temperature-independent and does not correlate with changes in egg weight during the incubation period.** (A-D) Phenotypic characteristics, differentiating male (A, B) and female (C, D) embryos. Arrow – presence of hemipenes in males (A) and absence in females (C); arrowhead – presence of heel spur on hind limbs of males (B) and absence in females (D). (E) Embryo sex distribution across different incubation temperatures, determined phenotypically and through genotyping for known male markers 23. No significant difference was identified for any tested temperature (Chi squared). (F) Final egg weights at the time of embryo dissection. The plotted data represent mean and standard deviation. At 28ºC male eggs were significantly heavier than female eggs ($p \leq 0.0283$ T-test). (G) Precent weight change for eggs is similar between males (blue) and females (red). No significant difference between percent weight change between males and females, as determined by a t-test. Number of eggs analyzed: 192, 2 of them were non-viable.
