## Supplementary Table 1 for "*Chamaeleo calyptratus* (veiled chameleon) chromosome-scale genome assembly and annotation provides insights into the evolution of reptiles and developmental mechanisms"

| ***C. Calyptratus* chromosomes (This study)** | Dovetail HiRise™ Assembly Scaffold | *C. Calyptratus* Flow Sorting Peak (Tishakova et al., 2022) | Z-linked genes  (Pokorna et al., 2011) | 45 rDNA  (Rovatsos et al., 2017) | Interstitial Telomeric Repeats |
| --- | --- | --- | --- | --- | --- |
| **1** | 2 | CCA 1 | 2^nd^ largest | “1^st^ pair” (Rovatsos et al., 2017)  Chromosome 1 (Tishakova et al., 2022) | “Pericentric regions of the largest metacentric chromosome pair” (Rovatsos et al., 2015) |
| **2** | 1 | CCA 2 |  |  |  |
| **3** | 4 | CCA 3 |  |  |  |
| **4** | 5 | CCA 4 |  |  |  |
| **5** | 6 | CCA 5 |  |  |  |
| **6** | 3 | CCA 6 |  |  |  |
| **7** | 3 | CCA 7 |  |  |  |
| **8** | 3 | CCA 8 |  |  |  |
| **9** | 3 | CCA 9 |  |  |  |
| **10** | 3 | CCA 10-11 |  |  |  |
| **11** | 3 | CCA 10-11 |  |  |  |
| **12** | 3 | CCA 12 |  |  |  |

**Supplementary Table 1.** **Cross-reference of the naming conventions used in this study for chromosomes and scaffolds, as well as other studies.** For studies identifying gene location based on genomic FISH, gene’s location is identified in the table in accordance with assembled genome data.

POKORNA, M., GIOVANNOTTI, M., KRATOCHVIL, L., KASAI, F., TRIFONOV, V. A., O'BRIEN, P. C., CAPUTO, V., OLMO, E., FERGUSON-SMITH, M. A. & RENS, W. 2011. Strong conservation of the bird Z chromosome in reptilian genomes is revealed by comparative painting despite 275 million years divergence. *Chromosoma,* 120**,** 455-68.

ROVATSOS, M., ALTMANOVA, M., JOHNSON POKORNA, M., VELENSKY, P., SANCHEZ BACA, A. & KRATOCHVIL, L. 2017. Evolution of Karyotypes in Chameleons. *Genes (Basel),* 8.

ROVATSOS, M., KRATOCHVIL, L., ALTMANOVA, M. & JOHNSON POKORNA, M. 2015. Interstitial Telomeric Motifs in Squamate Reptiles: When the Exceptions Outnumber the Rule. *PLoS One,* 10**,** e0134985.

TISHAKOVA, K. V., PROKOPOV, D. Y., DAVLETSHINA, G. I., RUMYANTSEV, A. V., O'BRIEN, P. C. M., FERGUSON-SMITH, M. A., GIOVANNOTTI, M., LISACHOV, A. P. & TRIFONOV, V. A. 2022. Identification of Iguania Ancestral Syntenic Blocks and Putative Sex Chromosomes in the Veiled Chameleon (Chamaeleo calyptratus, Chamaeleonidae, Iguania). *Int J Mol Sci,* 23.
