## Supplementary Table 2 for "*Chamaeleo calyptratus* (veiled chameleon) chromosome-scale genome assembly and annotation provides insights into the evolution of reptiles and developmental mechanisms"

| Positively correlated | | Negatively correlated | | No correlation | |
| --- | --- | --- | --- | --- | --- |
| Sequence | Correlation coefficient | Sequence | Correlation coefficient | Sequence | Correlation coefficient |
| CAGCCC | 0.948018 | ATTTCA | -0.95159 | AGCATG | -0.00637 |
| GGGCTG | 0.948018 | TGAAAT | -0.95159 | CATGCT | -0.00637 |
| CCCAGG | 0.943651 | AAATGT | -0.95008 | ACGCAT | -0.00541 |
| CCTGGG | 0.943651 | ACATTT | -0.95008 | ATGCGT | -0.00541 |
| CCCAGC | 0.94211 | AACATT | -0.9491 | GTAGAC | -0.00507 |
| GCTGGG | 0.94211 | AATGTT | -0.9491 | GTCTAC | -0.00507 |
| GGAGCC | 0.94032 | CAAATA | -0.94766 | GACTGA | -0.00502 |
| GGCTCC | 0.94032 | TATTTG | -0.94766 | TCAGTC | -0.00502 |
| CTGGCC | 0.940233 | TATGAA | -0.94572 | ACGACA | -0.00346 |
| GGCCAG | 0.940233 | TTCATA | -0.94572 | TGTCGT | -0.00346 |
| CGCCAG | 0.939429 | ACAAAT | -0.94356 | CAACTG | 0.002229 |
| CTGGCG | 0.939429 | ATTTGT | -0.94356 | CAGTTG | 0.002229 |
| CCAGCG | 0.939071 | ATCAAA | -0.94325 | CAGTCA | 0.002406 |
| CGCTGG | 0.939071 | TTTGAT | -0.94325 | TGACTG | 0.002406 |
| GAGCCC | 0.938727 | ATGAAA | -0.94302 | GGAACA | 0.007096 |
| GGGCTC | 0.938727 | TTTCAT | -0.94302 | TGTTCC | 0.007096 |
| CCCCAG | 0.937075 | AAATCA | -0.94302 | CACAAC | 0.009082 |
| CTGGGG | 0.937075 | TGATTT | -0.94302 | GTTGTG | 0.009082 |
| CCAGGG | 0.934418 | ATTTGA | -0.94266 | ACCAGT | 0.010413 |
| CCCTGG | 0.934418 | TCAAAT | -0.94266 | ACTGGT | 0.010413 |

**Supplementary Table 2 | Analysis of 6-mer distribution across the genome, in correlation with GC content.** 10 sequences each which have positive, negative and no correlation with GC content across the genome. Example graphical representations available in Supplementary Figure S10.
