## Supplementary Table 4 for "*Chamaeleo calyptratus* (veiled chameleon) chromosome-scale genome assembly and annotation provides insights into the evolution of reptiles and developmental mechanisms"

| Marker order | Marker designation from Nielsen et al. (2018) | Chromosome | Start | Finish |
| --- | --- | --- | --- | --- |
| 1 | confirmed_male_specific_reads_12_679 | chr5 | 1440 | 1972 |
| 2 | confirmed_male_specific_reads_5_639 | chr5 | 48110 | 47531 |
| 3 | confirmed_male_specific_reads_6_1,037 | chr5 | 130723 | 131313 |
| 4 | confirmed_male_specific_reads_10_752 | chr5 | 158950 | 158375 |
| 5 | confirmed_male_specific_reads_9_589 | chr5 | 234749 | 234193 |
| 6 | confirmed_male_specific_reads_8_526 | chr5 | 330208 | 329692 |
| 7 | confirmed_male_specific_reads_11_593 | chr5 | 460672 | 461317 |
| 8 | confirmed_male_specific_reads_13_424 | chr5 | 460675 | 460068 |
| 9 | confirmed_male_specific_reads_2_745 | chr5 | 635298 | 635949 |
| 10 | confirmed_male_specific_reads_3_771 | chr5 | 635301 | 634705 |
| 11 | confirmed_male_specific_reads_7_1,079 | chr5 | 850264 | 849645 |
| 12 | confirmed_male_specific_reads_4_785 | chr5 | 4719987 | 4720534 |
| 13 | confirmed_male_specific_reads_1_817 | chr5 | 11142043 | 11142672 |

**Supplementary Table 4 | Locations of male-specific markers on chromosome 5.** Markers were originally identified through RAD-sequencing of male and female genomes^1^.

1	Nielsen, S. V., Banks, J. L., Diaz, R. E., Jr., Trainor, P. A. & Gamble, T. Dynamic sex chromosomes in Old World chameleons (Squamata: Chamaeleonidae). *Journal of evolutionary biology* **31**, 484-490 (2018).
